# The plastidial protein MRC promotes starch granule initiation in wheat leaves but delays B-type granule initiation in the endosperm

**DOI:** 10.1101/2022.10.07.511297

**Authors:** Jiawen Chen, Yi Chen, Alexander Watson-Lazowski, Erica Hawkins, J. Elaine Barclay, Brendan Fahy, Robin Denley Bowers, Kendall Corbin, Frederick J. Warren, Andreas Blennow, Cristobal Uauy, David Seung

**Affiliations:** John Innes Centre, Norwich Research Park, Norwich, NR4 7UH, UK; Quadram Institute, Norwich Research Park, Norwich, NR4 7UQ, UK; University of Copenhagen, Thorvaldsensvej 40, 1871 Frederiksberg, Copenhagen, Denmark; Harper Adams University, Newport, Shropshire, TF10 8NB, UK; Department of Horticulture, College of Agriculture, Food and Environment, University of Kentucky, Lexington, Kentucky, USA

## Abstract

The spatial and temporal patterns by which starch granules initiate vary greatly between species and organs, but molecular factors that contribute to these diverse patterns are poorly understood. We reveal distinct organ-specific roles of the MYOSIN-RESEMBLING CHLOROPLAST PROTEIN (MRC) in regulating granule initiation in the endosperm and leaves of wheat. We isolated three independent TILLING mutants of tetraploid wheat (*Triticum turgidum* cv. Kronos) with premature stop or missense mutations in the A-genome homeolog, which we showed to be the only active homeolog in tetraploid wheat due to a disruption of the B-genome homeolog. Wheat endosperm contains both large A-type granules initiated during early grain development, and small B-type granules that initiate about 10 – 15 days later. The *mrc* mutants had significantly smaller A-type granules and a higher relative volume of B-type granules in the endosperm than the wild type. Whereas B-type granules initiated 15 - 20 days post anthesis (dpa) in the wild-type, they appeared as early as 10 dpa in the *mrc-1* mutant, suggesting a role for MRC in suppressing B-type granule initiation during early grain development. By contrast, MRC promotes granule initiation in leaves: mutants carrying premature stop mutations in *MRC* had fewer granules per chloroplast than the wild type. These contrasting roles of MRC among wheat organs provide new insight into functional diversification of granule initiation proteins, and suggest that they may facilitate the diverse patterns of granule initiation observed across species and organs.

## Introduction

Starch is the major storage carbohydrate in leaves, seeds and storage organs of most plants. It is synthesised in plastids as insoluble granules composed of two glucose polymers: amylopectin and amylose. Amylopectin is a branched polymer with linear α-1,4-glucan chains and α-1,6-branch points, and forms the semi-crystalline starch granule matrix - typically constituting 70-90% *w/w* of starch (Smith and Zeeman, 2020). Amylose constitutes 10 - 30% *w/w* of starch and is composed primarily of long α-1,4-linked linear chains (Seung, 2020). The biosynthesis of the starch polymers is relatively well understood at the molecular level, and is generally conserved between leaf and storage starches (Smith and Zeeman, 2020). However, we are only beginning to understand how starch granule formation is initiated, and the factors underpinning the vast diversity in granule initiation patterns observed between different organs and species (Tetlow and Emes, 2017; Seung and Smith, 2019; Chen et al., 2021).

A prime example of diverse granule initiation patterns between species can be observed in the seed endosperms of grasses (Matsushima et al., 2013). Grass species of the Triticeae, including important cereal crops such as wheat, barley and rye, have a unique bimodal size distribution of starch granules in the grain endosperm – containing large flattened A-type granules (20-30 μm in diameter) and small round B-type granules (2-7 μm in diameter) (Howard et al., 2011). The initiation of these two different types of granules is both spatially and temporally separated: A-type granules initiate in amyloplasts as early as 4 days post anthesis (dpa), whereas B-type granules initiate 10-15 days after the A-type granules (i.e., around 15-20 dpa) and at least partly within stromules that emanate from the amyloplast (Parker, 1985; Bechtel et al., 1990; Langeveld et al., 2000; Howard et al., 2011). This is distinct from most other grass species, which produce “compound” starch granules – where multiple granules initiate early during grain development in each amyloplast and eventually fuse (e.g., in rice) (Matsushima et al., 2013). Our recent work has identified proteins that are important for determining the initiation of bimodal starch granules in wheat. These include B-GRANULE CONTENT1 (BGC1) in *wheat/Aegilops* - which is orthologous to FLOURY ENDOSPERM6 (FLO6) in barley and rice and PROTEIN TARGETING TO STARCH 2 (PTST2) in Arabidopsis and *Brachypodium* (Peng et al., 2014; Saito et al., 2017; Seung et al., 2017; Chia et al., 2020; Watson-Lazowski et al., 2022). BGC1 in wheat has a dose-dependent effect on granule initiation, where partial reductions in gene dosage can almost eliminate B-type granules without affecting A-type granule formation, whereas complete loss of function also causes defective A-type granule formation, including the formation of some compound/semi-compound granules that arise from multiple initiations (Howard et al., 2011; Chia et al., 2020; Saccomanno et al., 2022). STARCH SYNTHASE 4 (SS4) is also required for normal A-type granule formation. In its absence, compound granules form in place of most A-type granules (Hawkins et al., 2021). The increased number of initiations per amyloplast that led to compound granule formation in these mutants was unexpected since in Arabidopsis leaves both SS4 and PTST2 promote granule initiation, and mutants lacking either protein have reduced numbers of starch granules per chloroplast (Roldán et al., 2007; Seung et al., 2017). These observations suggest that the proteins involved in granule initiation are to some extent conserved between species and organs, but they can act differently depending on the patterns of granule initiation in the species/tissue.

To gain further insight into such differences, we explored the function of wheat MYOSIN-RESEMBLING CHLOROPLAST PROTEIN (MRC, also known as PROTEIN INVOLVED IN STARCH INITIATION, PII1) in both chloroplasts of leaves and amyloplasts of the endosperm. MRC is a long coiled-coil protein that promotes granule initiation in Arabidopsis leaves, as most chloroplasts in the *mrc* mutant contain only a single granule (Seung et al., 2018; Vandromme et al., 2019). The exact function of MRC is unknown, but it is proposed to act via an interaction with SS4 and PTST2 (Seung et al., 2018; Vandromme et al., 2019). Consistent with previous work in Arabidopsis, we found that MRC promotes starch granule initiation in wheat leaves, but we discovered an unexpected, distinct role for MRC in the temporal control of B-type granule initiation in the wheat endosperm. MRC is expressed at the early stages of grain development, and wheat *mrc* mutants have severe alterations in the starch granule size distribution relative to the wild type, with smaller A-type granules and a higher relative volume of B-type granules. We demonstrate that this phenotype arises from the early initiation of B-type granules in the mutant, suggesting MRC represses B-type granule initiation during early grain development. This role of MRC in the wheat endosperm demonstrates how the function of granule initiation proteins can be adapted to mediate specific patterns of granule initiation among different species/tissues.

## Results

### The wheat orthologs of MRC are encoded on chromosomes 6A and 6D

The starch granule initiation protein MRC is highly conserved among land plants (Seung et al., 2018). To determine the role of MRC in wheat, we searched the wheat genome for genes encoding MRC orthologs. We ran a BLASTp search using the amino acid sequence of Arabidopsis MRC (*At*MRC, At4g32190) against the protein sequences from both tetraploid durum wheat (Svevo v1.1) (Maccaferri et al., 2019) and hexaploid bread wheat (IWGSG Chinese Spring) (Appels et al., 2018) genomes on Ensembl Plants. For hexaploid wheat (*Triticum aestivum*), the two top protein hits were TraesCS6A02G180500.1 (encoded on chromosome 6A) and TraesCS6D02G164600.1 (encoded on chromosome 6D), which shared 95% sequence identity with each other and were predicted as homeologs on Ensembl. Both genes had a two-exon structure like the Arabidopsis gene (Seung et al., 2018), and were in syntenic positions on the A and D genomes (Figure 1). For tetraploid durum wheat (*Triticum turgidum*), the top protein hit was TRITD6Av1G081580.1 (encoded on chromosome 6A), which was identical in nucleotide and amino acid sequence to TraesCS6A02G180500.1. To determine whether these proteins were true orthologs of *At*MRC, we repeated the phylogenetic analyses of MRC homologs from our previous study (Seung et al., 2018) with the wheat protein sequences included. The 6A and 6D proteins clustered together on the tree, distinctly within the grass clade containing the rice and maize sequences (Supplemental Figure 1A). This confirms that the proteins are the wheat orthologs of MRC. They will hereafter be referred to as *Ta*MRC-A1 (TraesCS6A02G180500) or *Tt*MRC-A1 (TRITD6Av1G081580), and *Ta*MRC-D1 (TraesCS6D02G164600). Notably, we did not find a full gene model for MRC on chromosome 6B, or anywhere else on the B genome, either in the durum or bread wheat genome.

**Figure 1:**
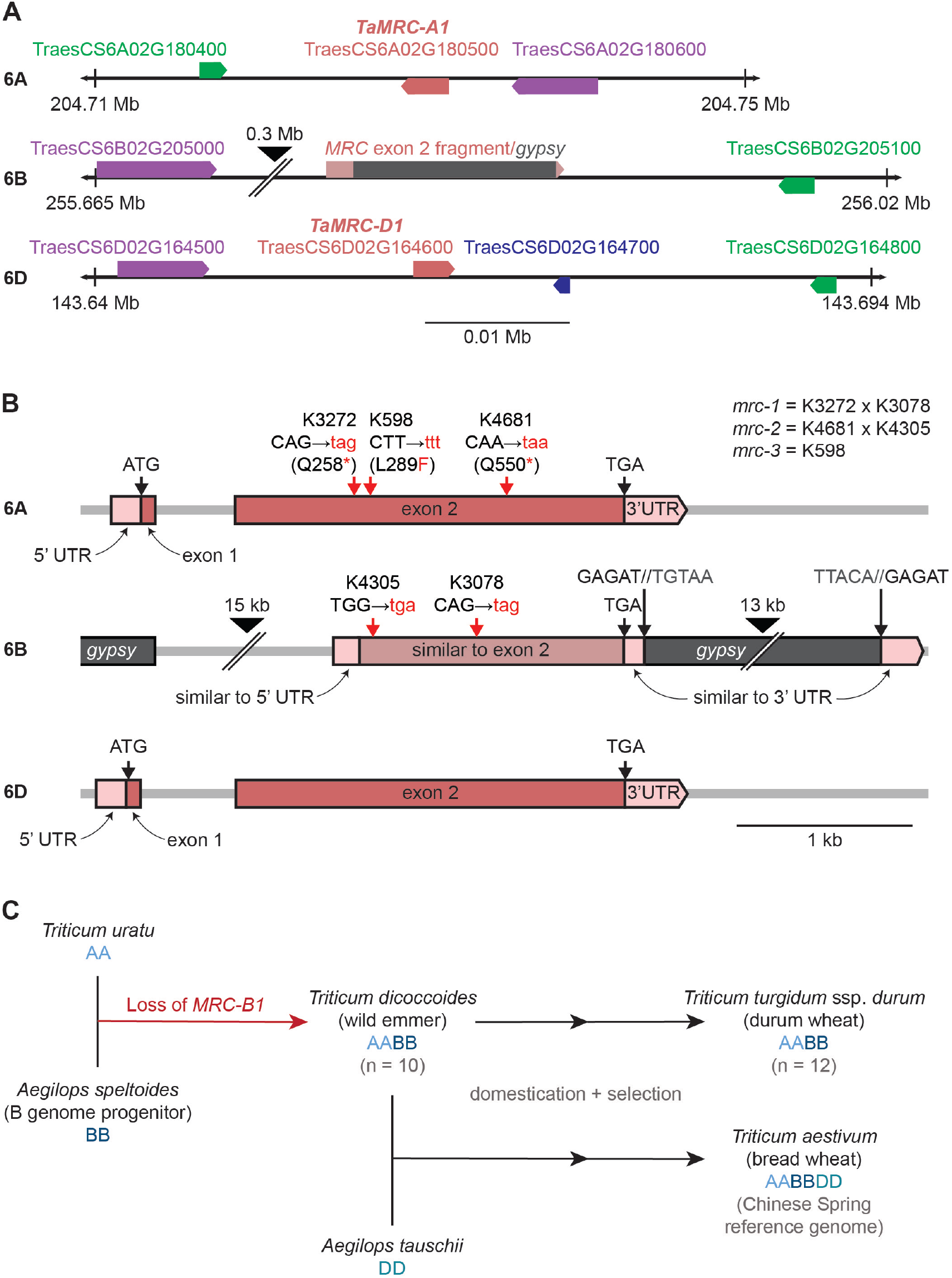
*MRC* homeologs in wheat are encoded on chromosomes 6A and 6D, with a disruption of the 6B homeolog. **A)** Location of *TaMRC* homeologs on chromosome 6A and 6D. The pink boxes represent *TaMRC* homeologs, while homeologs of the adjacent genes are shown in green (cytochrome P450 family protein), purple (respiratory burst oxidase homolog) and blue (uncharacterised protein). Arrowheads on the boxes indicate direction of transcription. The syntenic region on chromosome 6B has a large insertion, depicted with a black arrowhead. The diagram is drawn to scale, and chromosome coordinates of the region are indicated. **B)** Gene models of the *TaMRC-A1* and –*D1* homeologs and 6B pseudogene. Exons are represented with pink boxes, while light pink boxes represent the 5’ and 3’ UTRs. On the 6B region, areas with sequence similarity to exon 2 and UTRs of *TaMRC-A1* are indicated, as well as the location of *gypsy* retrotransposons. The locations of the mutations in the *mrc* mutants are depicted with red arrows, and the mutated codons/amino acids are shown in red letters. Large black arrowheads show where the sequence has been truncated for illustration - the length of truncated sequence is indicated above. **C)** Summary of species analysed for the loss of *MRC-B1* during wheat hybridisation

To investigate why no homeolog was detected on chromosome 6B, we looked at the syntenic region of chromosome 6B in Chinese Spring, where there was a stretch of sequence that had homology to exon 2 and the beginning of the 3’UTR (Figure 1A). Interestingly, around 14kb downstream of that, there was a region highly similar to the end of the 3’ UTR of *TaMRC-A1*. We looked at the transposable element annotation of the wheat reference genome around the exon 2 fragment (Daron et al., 2014), and a complete *gypsy* retrotransposon was annotated between the two 3’UTR fragments (Figure 1B). Further, we identified the 5 bp target site duplication (GAGAT, which is part of the 3’UTR) and the inverted terminal repeat (TGTAA and TTACA at the start and end of the retrotransposon, respectively) characteristic of retrotransposon insertions. The distance between the 5’ end of the exon 2 fragment and its upstream neighbouring gene (TraesCS6B02G20500, a respiratory burst oxidase homolog) was much larger than the distance between *Ta*MRC and the homeologs of the same neighbouring gene on 6A and 6D, indicating a large insertion in this 6B region (Figure 1A). Indeed, a fragment of another *gypsy* retrotransposon was found ca. 16 kb upstream of the exon 2 fragment. Additionally, there was a sequence with homology to the 5’ UTR of *TaMRC-A1* (85% identity over 243 bp) just 3 bp upstream of the exon 2 fragment, which suggested that a ~1.3 kbp deletion (based on A-genome distances) removed some of the 5’ UTR, all of exon 1, intron 1 and the start of exon 2 of *TaMRC-B1*. Similar to Chinese Spring, we found identical disruptions in *TaMRC-B1* sequences with retrotransposon insertions and deletions in ten additional wheat genome assemblies (Walkowiak et al., 2020). Overall, it appears that a deletion and a series of transposon insertions severely disrupted *MRC* on chromosome 6B in bread wheat.

Since a B-genome copy was also absent from durum wheat, it is likely that the disruption of *MRC* on chromosome 6B preceded the second hybridisation that resulted in hexaploid wheat. To further investigate when the disruption of *MRC-B1* occurred, we looked for *MRC-B1* in more tetraploid wheat accessions by aligning genome sequencing reads from *Triticum dicoccoides* (wild emmer) (n=10) and *Triticum turgidum* ssp. *durum* (pasta wheat) (n=12) against the A and B genomes of Chinese Spring (Zhou et al., 2020) (Supplemental Table 1). The exon 1 deletion and the retrotransposon insertion at the 3’ end were detected in all accessions (except for a few lines that had poor sequencing depth in the region), suggesting that *TaMRC* on 6B was disrupted before or immediately after the hybridization of diploid ancestors carrying the A and B genomes. To distinguish these possibilities, we examined *MRC* in *Aegilops speltoides*, the diploid species thought to be most closely related to the progenitor species of the wheat B genome. We ran a BLASTn search on the genome assembly of the *Aegilops speltoides* accession TS01 using the coding sequence of *TaMRC-A1* (Li et al., 2022) (Supplemental File 1). The top hit was on chromosome 6S (homologous to chromosome 6B in wheat) with intact exons one and two (97.5% sequence identity), and the translated protein sequence had 96% identity to *Ta*MRC-A1 with BLASTp. Thus, it is likely that *MRC* is intact in *Aegilops speltoides*, suggesting that the loss of the B-homeolog occurred shortly after the hybridisation that gave rise to tetraploid wheat (Figure 1C). It is therefore expected that all tetraploid wheats have one *MRC* homeolog (on chromosome 6A), and all hexaploids have two (on chromosomes 6A and 6D).

### MRC is expressed in leaves and developing endosperm

Using the hexaploid wheat expression browser (Borrill et al., 2016; Ramirez-Gonzalez et al., 2018), we found that transcripts of the 6A and 6D homeologs were present in both leaves and grains, suggesting that MRC plays a role in both these tissues (Figure 2A). To obtain temporal information on MRC expression in the endosperm during grain development, we performed RNAseq analysis on dissected endosperms of durum wheat (*Triticum turgidum* cv. Kronos) throughout grain development. The full dataset is available in our accompanying paper (Chen et al., 2022). The Kronos variety was chosen as it is the genetic background of the tetraploid wheat TILLING mutants characterised below. *TtMRC-A1* (TRITD6Av1G081580.1) showed a peak of expression during early grain development (8 dpa), but strongly decreased in expression between 10 – 20 dpa (Figure 2B). It therefore appears that *MRC* is expressed in the wheat endosperm almost exclusively during early grain development.

**Figure 2:**
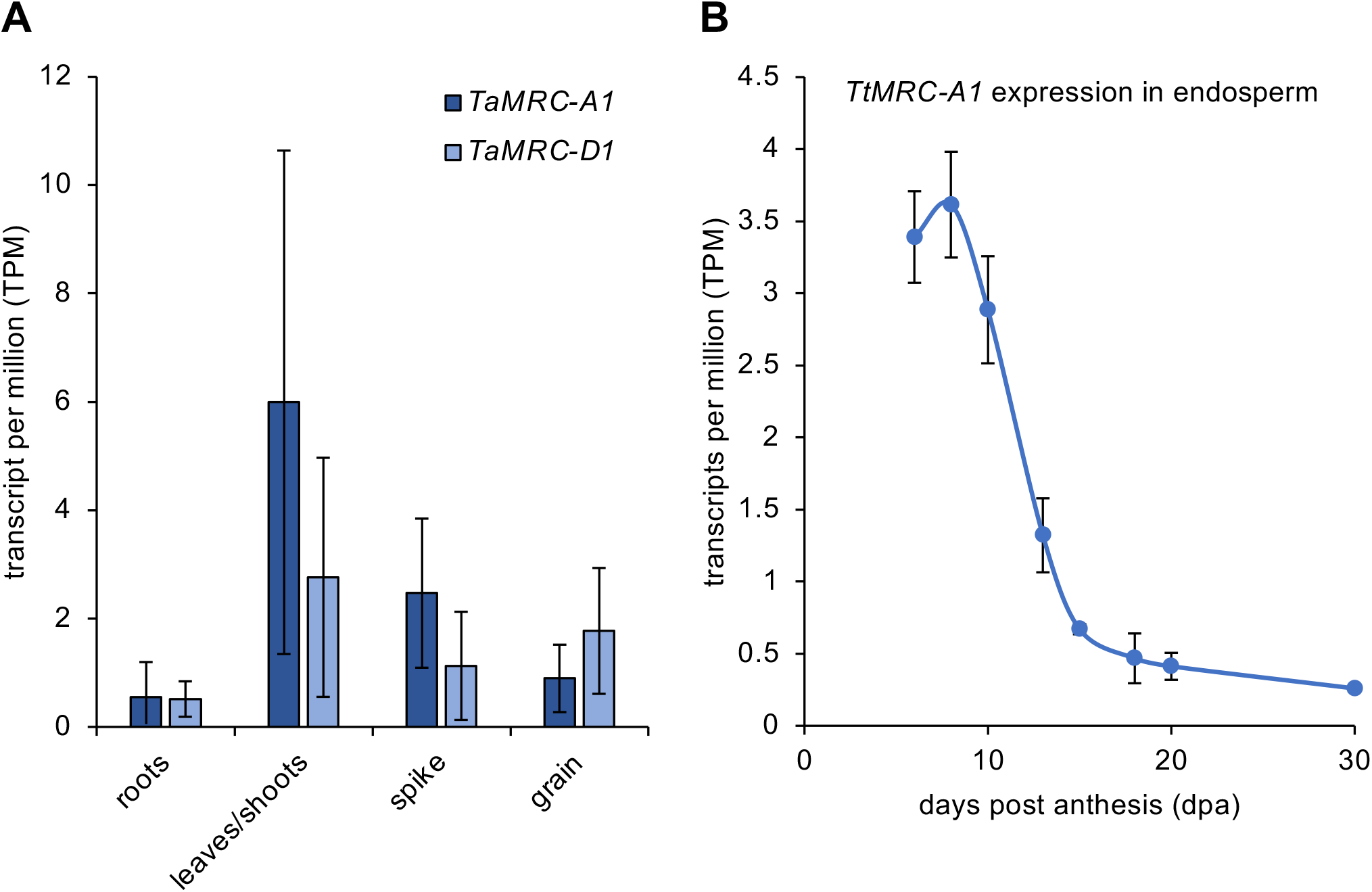
*MRC* is expressed in wheat leaves and wheat endosperm during early grain development. **A)** Expression of *TaMRC* homeologs between between different tissue types in *Triticum aestivum*. Data were obtained from the wheat expression browser, and values represent transcript per million (TPM) ± SEM from *n* = 89 (roots), 48 (leaves/shoots), 280 (spike), 166 (grain) samples from different experiments. **B)** Expression of *TtMRC-A1* (TRITD6Av1G081580.1) in the starchy endosperm during grain development of *Triticum turgidum* (variety Kronos). Data are from an RNAseq experiment described in Chen *et al*.(2022). Values represent TPM ± SEM from *n* = 3 for all timepoints.

### Loss of MRC does not affect the growth of wheat plants, or grain development

To study the function of MRC in wheat, we obtained mutants in durum wheat (*Triticum turgidum* cv. Kronos) defective in *MRC*. We used the wheat *in silico* TILLING mutant resource, which has an EMS-mutagenised population of Kronos with exome-capture sequencing data for identification of lines with mutations of interest (Krasileva et al., 2017). We obtained three mutants that were likely to cause a loss of function in *TtMRC-6A* (Figure 1B). The K3272 and K4681 lines contained premature stop codons in place of codons for the 258^th^ and 550^th^ amino acids respectively. In addition, we obtained a third line that contained a missense Leu289Phe mutation, which was predicted to be deleterious to protein function by SIFT scoring (Ng and Henikoff, 2006). The Leu^289^ residue is highly conserved in all MRC orthologs, and its mutation to a Phe residue is predicted to disrupt coiled coil formation in the region of the residue (Supplemental Figure 1B). Since the 6B homeolog of MRC has likely become a pseudogene, we predicted that *TtMRC-A1* would be the only functional *MRC* homeolog in tetraploid wheat. However, to rule out the possibility that the fragment of exon 2 on chromosome 6B affects MRC function, we also obtained the K4305 and K3078 lines which contain two different premature stop codon mutations in the putative reading frame of the exon. We generated the *mrc-1* lines by crossing K3272 and K3078, and the *mrc-2* lines by crossing K4681 and K4305, where we isolated lines homozygous for either the 6A (F_2_ *aa*BB) or 6B (F_2_ AA*bb*) mutation, or both (F_2_ *aabb*) The *mrc-1* double mutant line was backcrossed twice to wild type (WT), and the wild-type segregant (BC2F2 AABB) and the homozygous double mutant (BC_2_F_2_ *aabb*) were selected. These will be hereafter referred to as BC2 AABB and BC2 *aabb*. No backcrossing was done for the *mrc-2* line. The *mrc-3* line contained the K598 missense mutation, and no crossing was done.

There was no consistent effect of the independent *mrc* mutations on plant growth or grain development under our growth conditions. None of the *mrc* mutant plants appeared different from WT with respect to growth or the number of tillers per plant (Figure 3A, B, Supplemental Table 2A). The number of grains per plant also did not differ in the mutants, except for a slight decrease in *mrc-3* and the wild-type segregant (*mrc-1* BC2 AABB) compared to the WT (Figure 3C, Supplemental Table 2B). The morphology of the mature grains of the mutants was indistinguishable from the WT (Figure 3D), and there were no differences in thousand grain weight (TGW) and grain size between the WT and any of the three mutants *mrc-1, mrc-2, mrc-3*, or between the WT and the backcrossed *mrc-1* (*mrc-1* BC2 *aabb*); but the wild-type segregant (*mrc-1* BC2 AABB) had a slightly higher TGW and grain size compared to WT (Figure 3E, F, Supplemental Table 2C,D). This suggests that some of the background mutations in the wild-type segregant may have affected grain development, but these effects are small.

**Figure 3:**
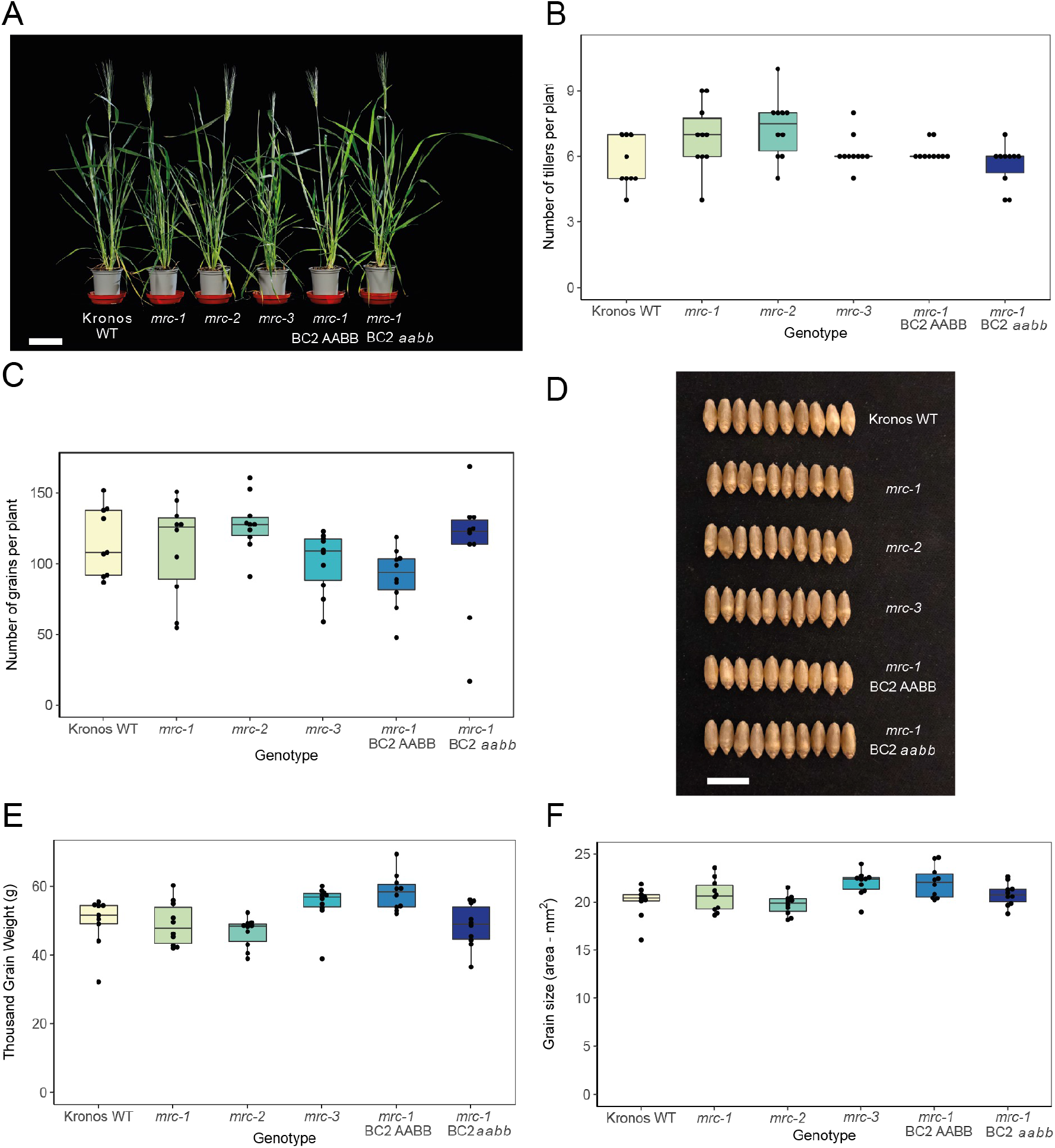
Mutations in *MRC* do not affect plant growth or the number and size of grains. **A)** Photographs of 7-week-old wild type (WT) and *mrc* mutant plants. Bar = 9 cm. **B)** Number of tillers per plant. **C)** Number of grains per plant. **D)** Photographs of mature grains. Bar = 1 cm. **E)** Thousand Grain Weight (TGW) of mature grains. **F)** Grain size, measured as the average area of individual mature grains from each plant. For statistical analysis of B, C, E, F, see Supplemental Table 1. For all boxplots, each box encloses the middle 50% of the distribution, the middle line is the median and the whiskers are the minimum and maximum values within 1.5 of the interquartile range. Dots are measurements from individual plants, with the same group of plants measured for all phenotypes and *n* = 9 – 10 individual plants.

### Loss of MRC greatly alters starch granule size distributions in the endosperm

To determine the effect of the *mrc* mutations on starch synthesis in grains, we first measured total starch content of mature grains. Starch content was largely similar between WT and the *mrc* mutants. Although some pairwise comparisons showed p<0.05 (WT with *mrc-2* and *mrc-3*), the confidence intervals of the difference in means was still close to zero, suggesting that any effect of the mutations was small (Figure 4A). Coulter counter measurements revealed that there were some minor differences between genotypes in the total number of granules relative to grain weight (granules/mg grain) (Figure 4B) but overall there was not a strong effect of loss of MRC. The backcrossed *mrc-1* mutant had more starch granules per unit grain weight than both the WT and the wild-type segregant, but the non-backcrossed *mrc-1* mutant was not significantly different to the WT. The *mrc-2* mutant also had relatively more granules than the WT, but *mrc-3* did not.

**Figure 4:**
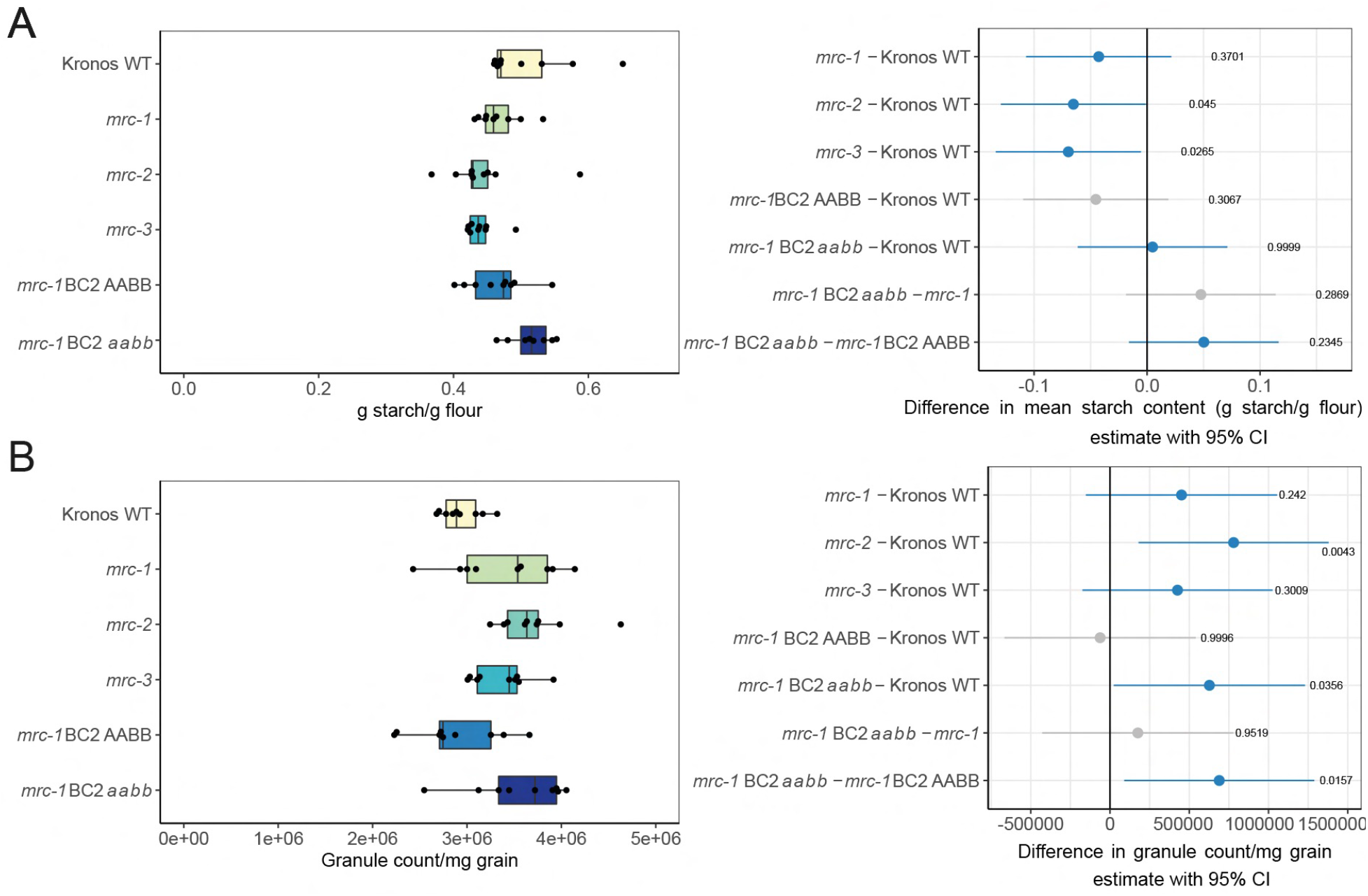
Mutations in *MRC* have variable effects on total starch in mature grains. A) Total endosperm starch content in mature grains in WT and *mrc* mutants. B) Number of starch granules per mg of grain, counted with a Coulter counter. For A and B, dots on the boxplots indicate individual plants, with *n* = 8 - 9. Each box encloses the middle 50% of the distribution, the middle line is the median and the whiskers are the minimum and maximum values within 1.5 of the interquartile range. All statistical analyses were performed using a linear model, with a one-way ANOVA and Tukey post-hoc test. Panels on the right indicate differences in means between genotypes from pairwise comparisons based on these models. The difference in means is indicated by a dot, with whiskers showing the 95% confidence interval (CI) of this difference, with the corresponding p-value. Grey indicates the WT or *mrc-1* mutant compared to the backcrossed line with the equivalent genotype at the *MRC* loci, and blue indicates all other pairwise comparisons.

Granule size distributions were determined from the Coulter counter data by plotting the percentage of starch volume in each size bin relative to the total volume of starch measured. We observed clear bimodal distributions for all genotypes, with a peak corresponding to A-type granules (~ 18 – 25 μm) and a peak corresponding to B-type granules (~3 – 10 μm). However, the distribution profiles in the mutants were very different from the WT, with the peak corresponding to B-type granules being more prominent in the mutants (Figure 5A). The profiles of the backcrossed *mrc-1* lines looked similar to their non-backcrossed equivalents. We fitted a bimodal log-normal distribution curve to the profiles of each sample to estimate the total volume percentage of B-type granules and the mean sizes of A- and B-type granules. Comparing the means of these extracted values between genotypes showed a higher B-type granule percentage (by volume) for all three mutants (*mrc-1, mrc-2* and *mrc-3*) compared to the WT (Figure 5B). The strongest increase was seen for *mrc-1*, and this increase was consistent when comparing the double backcrossed *mrc-1 (mrc-1* BC2 *aabb*) with WT and the wild-type segregant (*mrc-1* BC2 AABB). There was a small increase in the volume percentage of B-type granules in the wild-type segregant compared to the WT, but the difference was much smaller than between the other genotypes. The higher B-type granule volume percentage in *mrc* mutants could be due to an increase in both B-type granule size and number. The mean B-type granule size was larger than WT for *mrc-1* (and *mrc-1* BC2 *aabb*) and *mrc-2*, but not *mrc-3* (Figure 5D). By contrast, the mean A-type granule diameter was smaller for all three mutants than for WT (Figure 5C). Thus, the increased proportion of B-type granule volume may be due to a combination of smaller A-type granules and larger and/or more numerous B-type granules.

**Figure 5:**
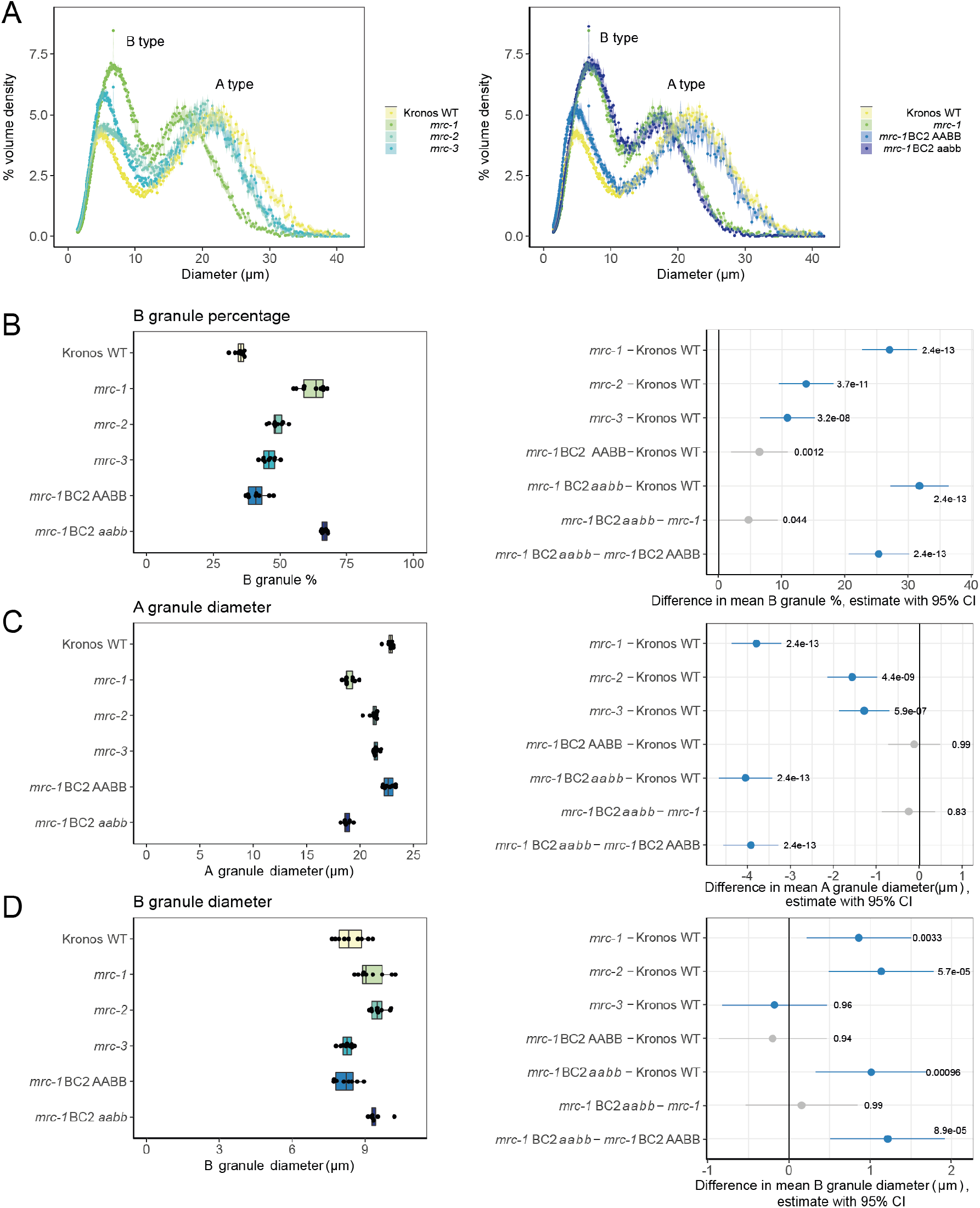
Endosperm starch in mature grains of *mrc* mutants have altered granule size distribution. **A)** Coulter counter traces with evenly binned x-axes show a bimodal distribution of granule sizes from purified wheat endosperm starch. Data points are mean values from 9 individual plants of each genotype (3 grains from each plant), with the standard error of the mean shown as a shaded ribbon. WT and *mrc-1* in the left and right panel are the same data. **B, C, D)** Mean B-type granule percentage volume (B), A-type granule diameter (C) and B-type granule diameter (D) values extracted from bimodal log-normal distribution curves fitted to Coulter counter traces of individual plants. Extracted values and boxplots are shown on the left, where dots indicate mean values from individual plants (*n* = 9). Each box encloses the middle 50% of the distribution, the middle line is the median and the whiskers are the minimum and maximum values within 1.5 of the interquartile range. All statistical analyses were performed using linear models for each panel, with a one-way ANOVA and Tukey post-hoc test. Panels on the right indicate differences in means between genotypes from pairwise comparisons based on these models. The difference in means is indicated by a dot, with whiskers showing the 95% confidence interval (CI) of this difference, with the corresponding p-value. Grey indicates the WT or *mrc-1* mutant compared to the backcrossed line with the equivalent genotype at the *MRC* loci, and blue indicates all other pairwise comparisons.

We also explored whether the *mrc* mutations affected starch granule shape. Examination of iodine-stained thin sections of mature grains using light microscopy showed that, like the WT, all mutants had flattened A-type and round B-type granules (Figure 6A). Similarly, no defects in A- or B-type granule shape were observed in the mutants using scanning electron microscopy (SEM) (Figure 6B). Starch polymer structure and granule composition was not affected by loss of functional MRC. The *mrc-1* mutant, with the strongest alteration in granule size distribution, had normal amylopectin structure and amylose content (Supplemental Figure 2).

**Figure 6:**
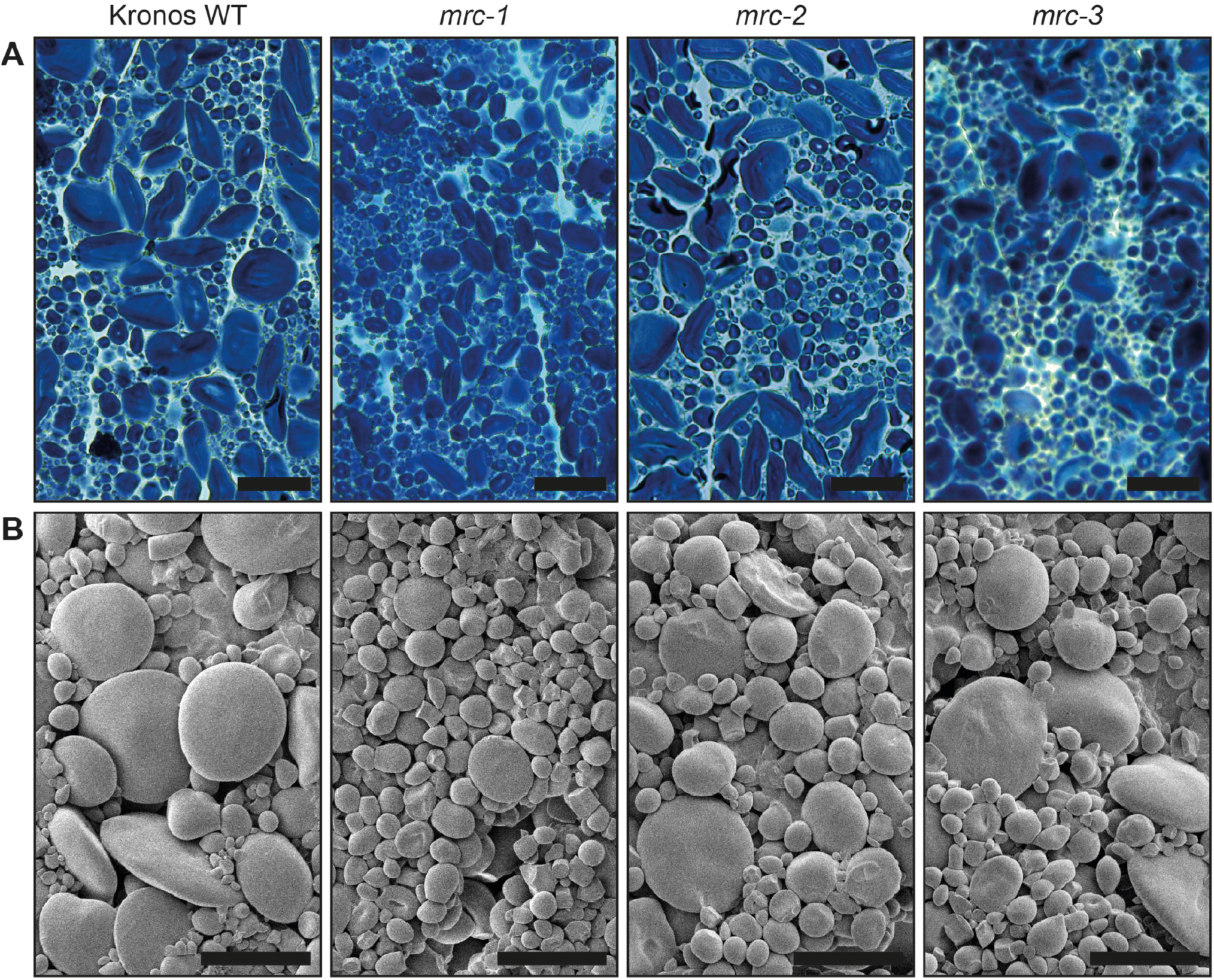
Endosperm starch granules of mature grains in three wheat *mrc* mutants are similar in shape compared to wild type. **A)** Thin sections of mature endosperm tissue were stained with Lugol’s solution and imaged using light microscopy. Bar = 40 μm. **B)** Purified endosperm starch granules were observed using scanning electron microscopy (SEM). Bar = 20 μm.

We sought experimental evidence about the contribution of the fragment of exon 2 on chromosome 6B to the observed differences in granule size distribution between genotypes (Supplemental Figure 3). Quantification of starch granule size distribution in the full set of homozygous genotypes resulting from the crosses that yielded the *mrc-1* and *mrc-2* mutants [indicated as *aaBB* (6A mutant), *AAbb* (6B mutant) and *aabb* (6A and 6B double mutant)] showed that WT and AA*bb* genotypes had identical granule distributions. The *aa*BB and *aabb* had different distributions from the WT but were identical to each other. These data showed that the fragment of exon 2 on chromosome 6B has no influence on granule size distribution. They are consistent with the suggestion that this is a pseudogene, and hence that the 6A copy of *MRC* is the only functional homeolog in tetraploid wheat.

Overall, these data suggest that MRC is required for the normal size distribution of starch granules in wheat endosperm. In tetraploid wheat, mutants lacking in the 6A copy of *MRC* consistently had a higher relative volume of B-type granules in the endosperm than the WT, and smaller A-type granules. This change in granule size distribution occurred without accompanying changes in total starch content, starch granule shape, amylose content or amylopectin structure.

### Loss of MRC results in the early initiation of B-type granules

To understand how MRC affects the size distribution of endosperm starch granules, and its specific effects on A-type or B-type granules, we investigated granule initiation during grain development in the *mrc* mutant with the strongest phenotype, *mrc-1*. We measured the total starch content and number of starch granules in dissected endosperms of developing grains harvested 8, 14, 20 and 30 days post anthesis (dpa). The total starch content of the endosperm increased between each time point, and there was no significant difference between the mutant and the WT at any time point (Figure 7A). At the 8 dpa timepoint, the mutant and the WT contained similar numbers of starch granules. Interestingly, for the two subsequent time points (14 dpa and 20 dpa), the mutant endosperms contained almost twice as many starch granules as the WT, despite similar starch contents (Figure 7B). The largest increase in granule number during grain filling was observed between the 20 and 30 dpa timepoints in the WT, and between the 14 and 20 dpa timepoints in the mutant. At the 30 dpa timepoint, the difference in granule number between the mutant and WT decreased. We also noted that in both the WT and mutant, the number of starch granules decreased between the 8 and 14 dpa timepoints. The reason for this is unknown, but it has also been observed in *Aegilops* species – which are close relatives of wheat (Howard et al., 2011).

**Figure 7:**
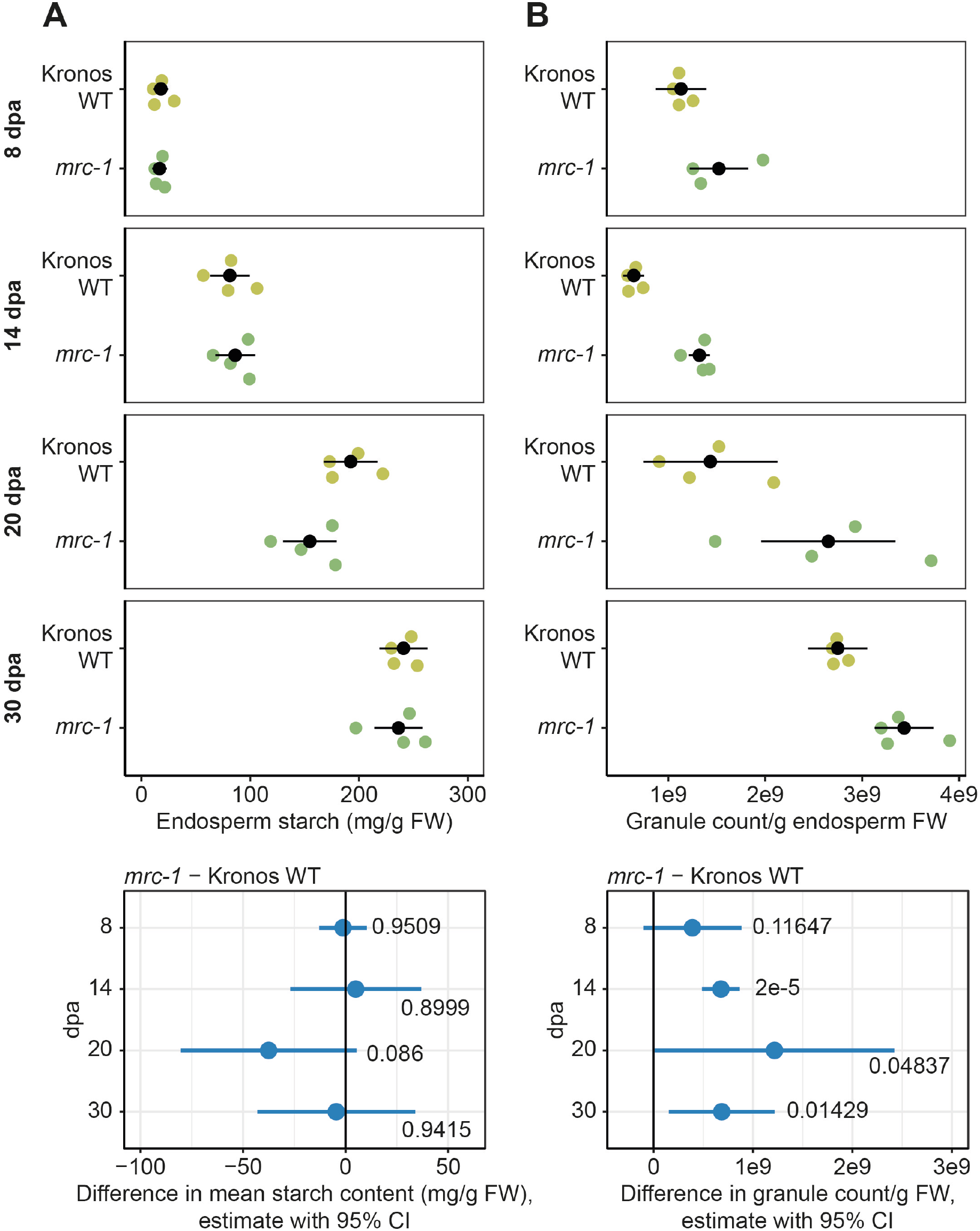
In developing endosperm, granule number increases in *mrc-1* compared to WT despite having similar starch content. The endosperm was dissected from developing grains of WT and *mrc-1*, harvested at 8, 14, 20 and 30 dpa, with *n* = 3 - 4 individual plants for each genotype per time point, indicated by coloured dots in the upper panels, with the mean ± 95% confidence interval (CI) in black dots and whiskers. **A)** Starch content of the endosperm. Values are expressed relative to the fresh weight of the dissected endosperm. **B)** Starch granule number in the endosperm. Starch was purified from dissected endosperm and the number of granules was determined using a Coulter counter. Values are expressed relative to the fresh weight of the dissected endosperm. For A and B, individual linear models were fitted to the data of each time point, with a one-way ANOVA and Tukey post-hoc test to compare the means of WT and *mrc-1*. Panels on the bottom summarise the differences in means from these linear models, indicated by a dot, with whiskers showing the 95% CI of this difference, with the corresponding p-value.

In WT endosperm, there was a unimodal distribution of starch granule sizes at the 8 and 14 dpa timepoints, and only A-type granules with their characteristic flattened morphology were observed using SEM (Figure 8). The A-type granules grew substantially in size between the two timepoints, seen as a shift in the granule size distribution peak. B-type granules only became prominent at the 20 dpa timepoint. In the *mrc-1* mutant A-type granules were initially the same size as those of WT (at 8 dpa), but subsequently grew more slowly than wild-type granules. By contrast, B-type granules were already present at 14 dpa in the *mrc-1* mutant, (seen as a distinct shoulder that appeared in the granule size distribution), considerably earlier than in the WT. Taken together, these data suggest that the larger number of granules observed between 14-20 dpa in the *mrc-1* endosperm compared to the WT (observed in Figure 7B) is due to the early initiation of B-type granules in the mutant.

**Figure 8:**
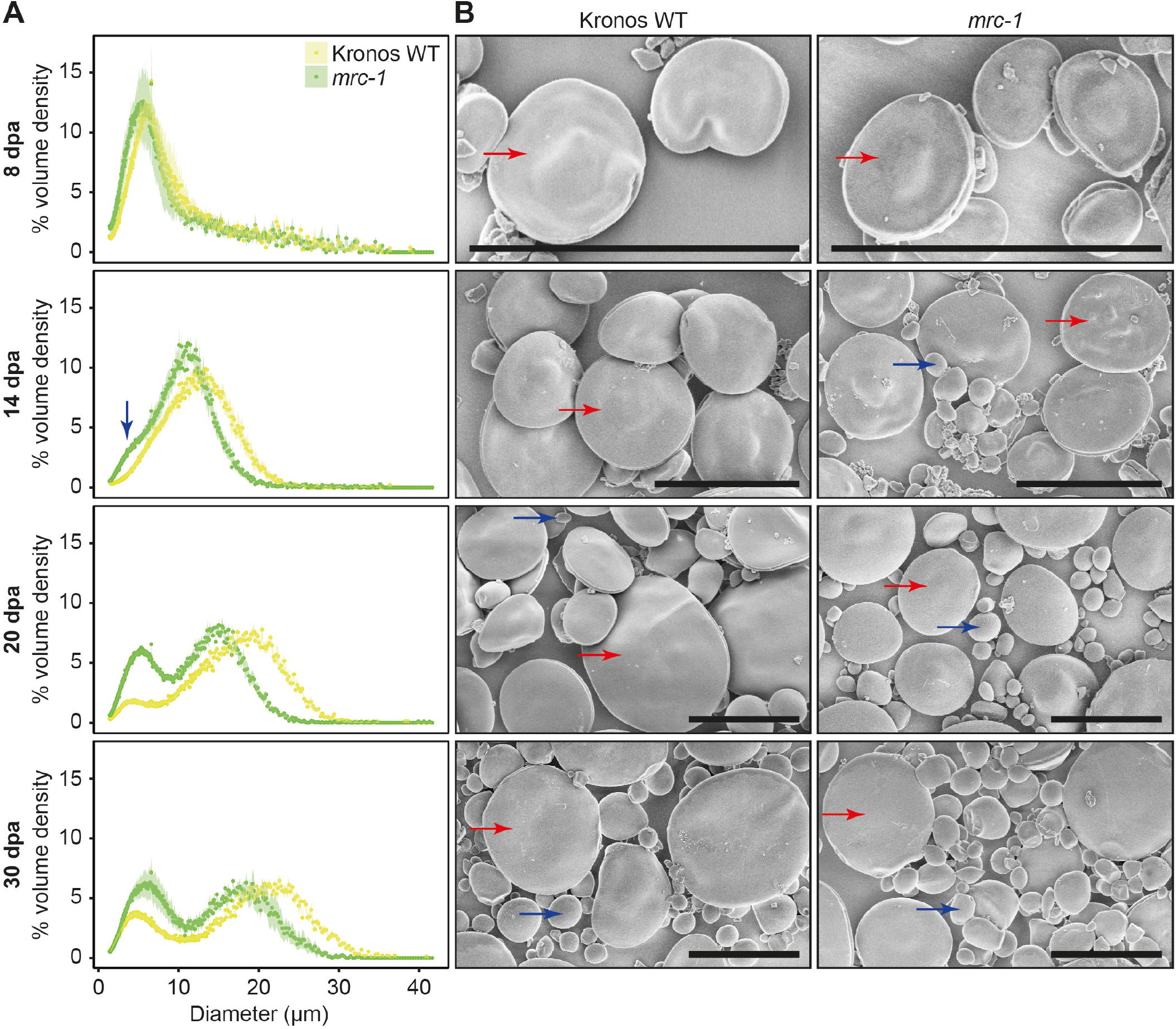
The *mrc-1* mutant initiates B-type granules earlier in grain development than in the wild type (WT). The endosperm was dissected from developing grains of WT and *mrc-1* harvested at 8, 14, 20 and 30 dpa. **A)** Coulter counter traces with evenly binned x-axes show the starch granule size distribution of endosperm starch. Distributions are the average of *n* = 4 measurements, each carried out on grains harvested from a different plant (three grains per measurement). Data points are mean values from these 4 measurements, with the standard error of the mean shown as a shaded ribbon. The B-type granule peak in the *mrc-1* mutant at 14 dpa is indicated with a blue arrow. **B)** Endosperm starch granules were observed using scanning electron microscopy (SEM). Bars = 20 μm. Examples of A-type granules and B-type granules are marked with red and blue arrows respectively.

B-type granules typically initiate in close proximity to each other–appearing as ‘clusters’ in between the A-type granules – and at least some B-type granules form in amyloplast stromules (Parker, 1985; Langeveld et al., 2000). Given the unusual timing of B-type granule initiation in *mrc-1*, we explored whether the loss of MRC also affected the location of B-type granule initiation. First, we harvested grains during their development (10, 15, and 20 dpa), subjected them to critical point drying, and imaged the cut face of sections through the endosperm using SEM. Consistent with the findings from the purified starch granules (Figure 8), B-type granules were already present at 10 dpa in the mutant, whereas they only became prominent after 20 dpa in the WT (Figure 9A). The B-type granules occurred in clusters in the mutant that resembled those of the WT. Secondly, we examined sections off developing grains using light and electron microscopy. For light microscopy, sections were stained with toluidine blue (a negative stain for starch). At 15 dpa, most starch granules in the wild-type endosperm were flattened A-type granules, and very few B-type granules were visible (Figure 9B). However, in the endosperm of the *mrc-1* mutant, many clusters of B-type granules were present. Using transmission electron microscopy (TEM), we investigated whether these clusters of B-type granules occurred within single amyloplasts, and in particular in stromules. Indeed, multiple B-type granules were enclosed within amyloplasts, and the elongated morphology of these amyloplast regions strongly suggested that they are stromules (Figure 9C). It is difficult to determine the exact percentage of B-type granules in stromules, because stromules are difficult to observe in two-dimensional sections. However, apart from their earlier occurrence in the mutant, we did not observe anything unusual about the location of B-type granules in the *mrc-1* mutant.

**Figure 9:**
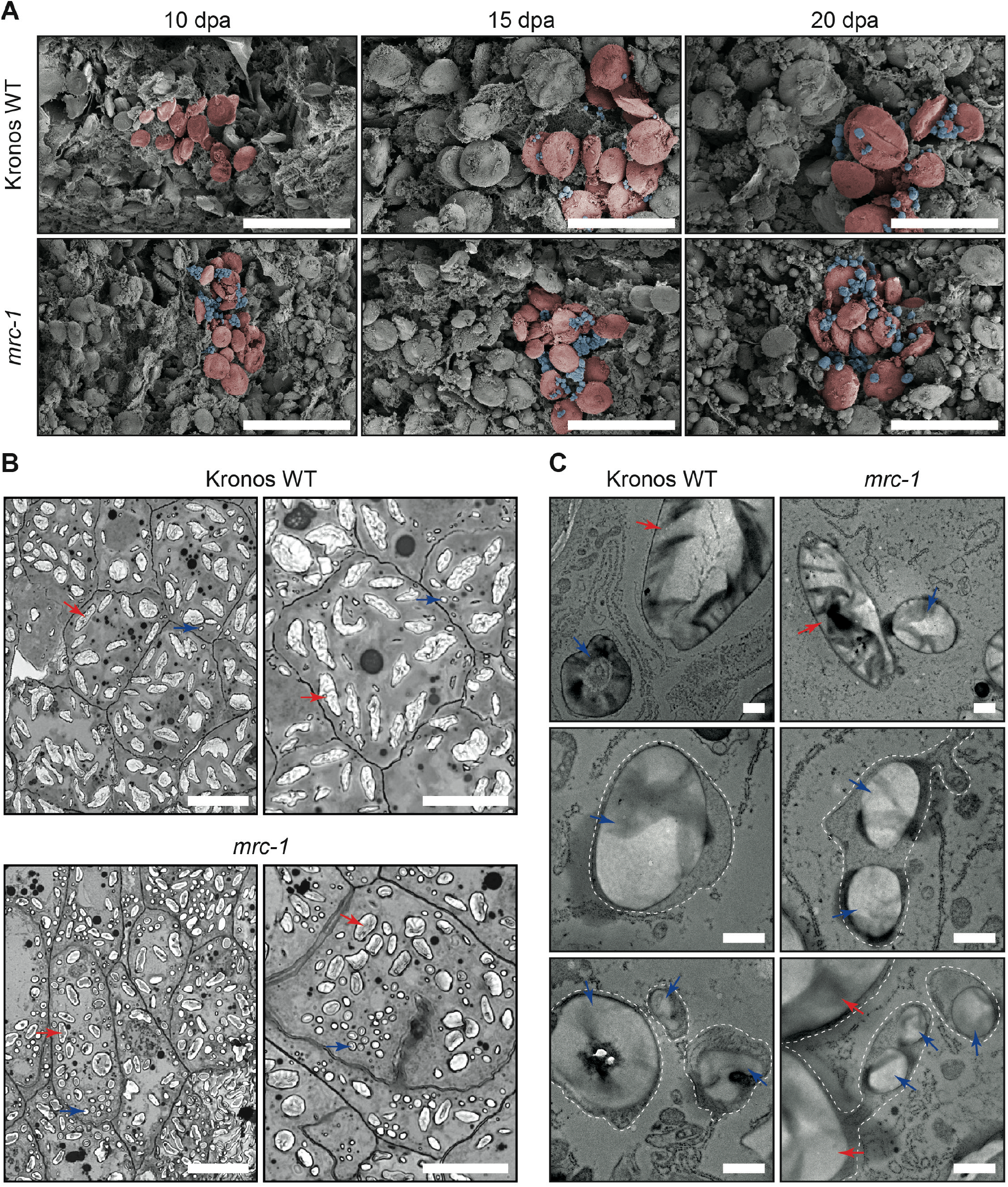
The early initiating B-type granules in *mrc-1* occur at least partially in stromules. **A)** Scanning electron micrographs of developing endosperm tissue subjected to critical point drying. Grains were harvested from WT and *mrc-1* at 10, 15 and 20 days post anthesis (dpa). A representative region of each panel has been pseudo-coloured, with A-type granules in red shading and B-type granules in blue shading. Bars = 50 μm. **B)** Light micrographs of endosperm sections. Semi-thin sections of embedded developing grains (15 dpa) were stained with toluidine blue as a negative stain for starch granules. Examples of A-type granules and B-type granules are marked with red and blue arrows respectively. Bars = 50 μm. **C)** Endosperm sections observed using transmission electron microscopy, using the same samples as in B. The periphery of amyloplast membranes, where visible, are indicated with a dotted white line. Bars = 1 μm.

In conclusion, MRC is required for the temporal control of B-type granule initiation during wheat grain development. It is expressed during early grain development, and its loss leads to the early initiation of B-type granules. We therefore propose that MRC acts as a repressor of B-type granule formation in the developing wheat endosperm during early grain development.

### TaMRC promotes granule initiation in wheat leaves

Considering the role of MRC in promoting granule initiation in Arabidopsis leaves (Seung et al., 2018; Vandromme et al., 2019), it was surprising that MRC repressed B-type granule formation in wheat endosperm. We therefore investigated whether this repressive role also applies to granule initiation in wheat leaves, which would suggest a divergence in MRC function between wheat and Arabidopsis; or whether the role of MRC in wheat leaves is the same as in Arabidopsis but it has a distinct function in granule initiation in the endosperm.

We found that *mrc-1* and *mrc-2* had fewer granules per chloroplast than the WT (Figure 10 A, C, Table 1). As in the endosperm, the *mrc-1* mutant showed the strongest effect, having almost 50% fewer granules per chloroplast than the WT (Table 1). Interestingly, there was no effect of *mrc-3*, suggesting that the L289F mutation does not affect MRC function in leaves (Figure 10 A, C, Table 1). To assess granule number, shape and size in leaf chloroplasts of our wheat mutants, leaf tissue was harvested from the middle of the older of two leaves of 10-day-old wheat seedlings and sections were imaged with light microscopy (Figure 10A, B). The number of granules per chloroplast was quantified from these images. We measured three individual plants per genotype for each experiment and compared the mean granules per chloroplast between genotypes using a negative binomial mixed effects model with individual biological replicates as random effect. We first compared all three *mrc* mutants with the WT (Figure 10C, Table 1), then we compared the backcrossed *mrc-1* lines with the WT (Figure 10D, Table 1).

**Figure 10:**
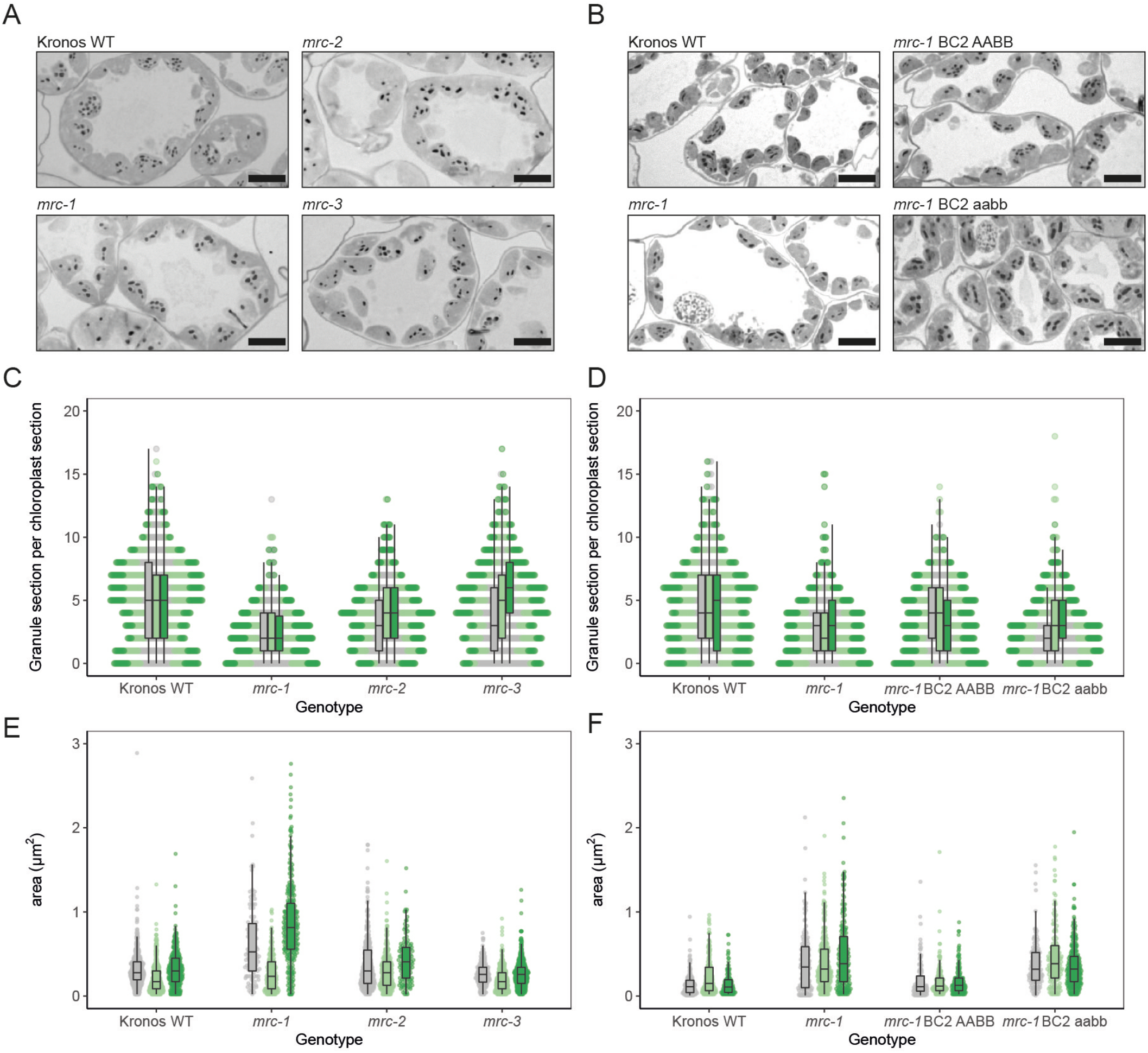
Chloroplasts of *mrc-1* mutants have fewer and larger granules than the wild type. **A, B)** Representative light microscopy images of thin sections (500 nm) from the middle of the older of two leaves in 10-day old wheat seedlings, collected at the end of day. Sections were stained for starch using periodic acid and Schiff staining. Scale bar = 10 μm. **C, D)** Distributions of counts of granule sections per chloroplast section, counted from light microscopy images as in A and B. The three different colours correspond to three individual plants of each genotype, per experiment. For each sample, 200 – 250 chloroplasts were counted, with absolute values varying between samples. Dots represent individual chloroplasts. **E, F)** Distributions of starch granule sizes measured as the area in the light microscopy images as in A and B. The three different colours correspond to three individual plants of each genotype, per experiment. 100 – 400 granules were measured in each sample. Dots represent individual granules. For all boxplots, each box encloses the middle 50% of the distribution, the middle line is the median and the whiskers are the minimum and maximum values within 1.5 of the interquartile range. Measurements were made on the same batch of plants as Figure 11, experiment 2 (panels A, C, E here) and experiment 5 (panels B, D, F here).

**Table 1.** Statistics of pairwise comparisons of leaf granule per chloroplast in young wheat leaves, from negative binomial mixed effect models with biological replicate (3 of each genotype) as random effect, using Tukey post-hoc tests. Individual models for each experiment were used. SE = standard error, 95% CI = 95% confidence interval.

| contrast | Experiment | ratio | SE | - 95% CI | + 95 CI | z.ratio | p.value |
| --- | --- | --- | --- | --- | --- | --- | --- |
| WT / *mrc-1* | 2 | 1.997 | 0.190 | 1.564 | 2.550 | 7.271 | 2.17E-12 |
| WT / *mrc-2* | 2 | 1.390 | 0.131 | 1.091 | 1.769 | 3.497 | 0.00265 |
| WT / *mrc-3* | 2 | 1.091 | 0.102 | 0.858 | 1.387 | 0.929 | 0.789273 |
| *mrc-1* / *mrc-2* | 2 | 0.696 | 0.067 | 0.544 | 0.890 | -3.783 | 0.000892 |
| *mrc-1* / *mrc-3* | 2 | 0.546 | 0.052 | 0.427 | 0.698 | -6.336 | 1.41E-09 |
| *mrc-2* / *mrc-3* | 2 | 0.785 | 0.074 | 0.616 | 1.000 | -2.564 | 0.05063 |
| *mrc-1* BC2 *aabb* / *mrc-1* | 5 | 1.030 | 0.097 | 0.810 | 1.311 | 0.318 | 0.988897 |
| *mrc-1* BC2 *AABB* / *mrc-1* | 5 | 1.234 | 0.115 | 0.971 | 1.569 | 2.256 | 0.108551 |
| *mrc-1* BC2 *AABB* / *mrc-1* BC2 *aabb* | 5 | 1.198 | 0.111 | 0.943 | 1.522 | 1.943 | 0.210022 |
| WT / *mrc-1* | 5 | 1.635 | 0.152 | 1.287 | 2.077 | 5.285 | 7.52E-07 |
| WT / *mrc-1* BC2 *aabb* | 5 | 1.587 | 0.147 | 1.251 | 2.014 | 4.982 | 3.76E-06 |
| WT / *mrc-1* BC2 *AABB* | 5 | 1.325 | 0.122 | 1.045 | 1.679 | 3.048 | 0.012347 |

The backcrossed *mrc-1* (*mrc-*1 BC2 *aabb*) had fewer granules per chloroplast than WT, similar to the non-backcrossed *mrc-1*. However, we were not able to make meaningful comparisons with the wild-type segregant (*mrc-*1 BC2 AABB) because – as for comparisons of TGW and grain size above – values for the wild-type segregant differed from those of the WT, with significantly fewer granules per chloroplast than WT. Overall, the smaller number of granules in *mrc* chloroplasts than in WT chloroplasts suggests that MRC promotes granule initiation in wheat leaves, pointing to distinct roles of MRC in different organs.

While the effect of the *mrc-1* mutation on starch granule number per chloroplast was consistent, two other aspects of leaf starch – granule size and total starch content – appeared to be interdependent and varied between our experiments. We quantified the size of granules from these images by measuring the area of granules in the sections. The distributions showed a trend towards larger granule sizes in the *mrc-1* mutant (Figure 10E, F). However, after fitting a linear mixed effects model with individual biological replicates as random effect (Table 2), pairwise comparisons showed no differences between any of the non-backcrossed *mrc* mutants and WT. However, in the experiment comparing the backcrossed *mrc-1* mutants to WT and wild-type segregant, the granules were larger in both backcrossed and non-backcrossed *mrc-1* mutants. Aside from the differences in size, there was no visible difference in starch granule shape in any of the mutants compared to WT.

**Table 2.**
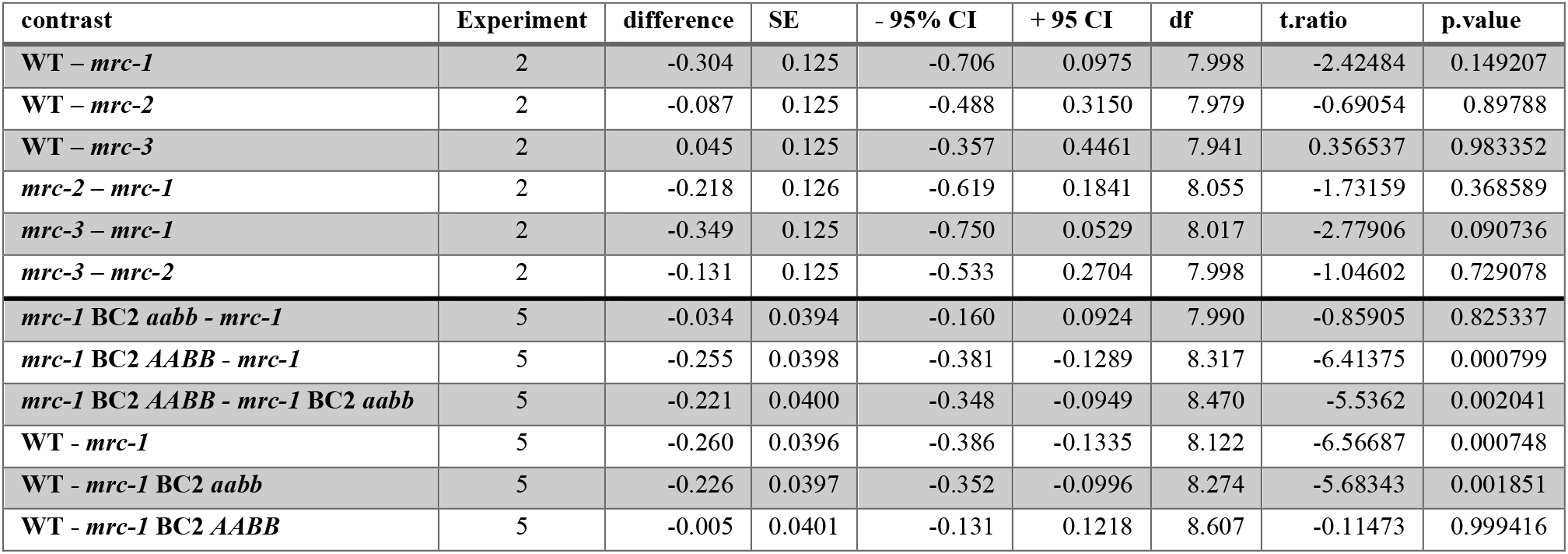
Statistics of pairwise comparisons of granule size in young wheat leaves, from linear mixed effect models with biological replicate (3 of each genotype) as random effect, using Tukey post-hoc tests and Satterthwaite degrees of freedom calculation. Individual models for each experiment were used. SE = standard error, df = degrees of freedom, 95% CI = 95% confidence interval.

| contrast | Experiment | difference | SE | - 95% CI | + 95 CI | df | t.ratio | p.value |
| --- | --- | --- | --- | --- | --- | --- | --- | --- |
| WT - *mrc-1* | 2 | -0.304 | 0.125 | -0.706 | 0.0975 | 7.998 | -2.42484 | 0.149207 |
| WT - *mrc-2* | 2 | -0.087 | 0.125 | -0.488 | 0.3150 | 7.979 | -0.69054 | 0.89788 |
| WT - *mrc-3* | 2 | 0.045 | 0.125 | -0.357 | 0.4461 | 7.941 | 0.356537 | 0.983352 |
| *mrc-2* - *mrc-1* | 2 | -0.218 | 0.126 | -0.619 | 0.1841 | 8.055 | -1.73159 | 0.368589 |
| *mrc-3* - *mrc-1* | 2 | -0.349 | 0.125 | -0.750 | 0.0529 | 8.017 | -2.77906 | 0.090736 |
| *mrc-3* - *mrc-2* | 2 | -0.131 | 0.125 | -0.533 | 0.2704 | 7.998 | -1.04602 | 0.729078 |
| *mrc-1* BC2 *aabb* - *mrc-1* | 5 | -0.034 | 0.0394 | -0.160 | 0.0924 | 7.990 | -0.85905 | 0.825337 |
| *mrc-1* BC2 *AABB* - *mrc-1* | 5 | -0.255 | 0.0398 | -0.381 | -0.1289 | 8.317 | -6.41375 | 0.000799 |
| *mrc-1* BC2 *AABB* - *mrc-1* BC2 *aabb* | 5 | -0.221 | 0.0400 | -0.348 | -0.0949 | 8.470 | -5.5362 | 0.002041 |
| WT - *mrc-1* | 5 | -0.260 | 0.0396 | -0.386 | -0.1335 | 8.122 | -6.56687 | 0.000748 |
| WT - *mrc-1* BC2 *aabb* | 5 | -0.226 | 0.0397 | -0.352 | -0.0996 | 8.274 | -5.68343 | 0.001851 |
| WT - *mrc-1* BC2 *AABB* | 5 | -0.005 | 0.0401 | -0.131 | 0.1218 | 8.607 | -0.11473 | 0.999416 |

Similar variability between experiments was seen in total starch content, which was measured in both leaves of 10-day-old wheat seedlings at the end of the day, when maximum starch is expected. Our first three experiments compared the genotypes WT, *mrc-1, mrc-2* and *mrc-3* (Experiments 1, 2, 3; Supplemental Table 4), and the pooled data showed a lower starch content in all *mrc* mutants compared to WT (Figure 11A). Our next two experiments (Experiments 4 and 5, Supplemental Table 4) compared end of day starch content in WT, *mrc-1, mrc-1* BC2 *aabb* and *mrc-1* BC2 AABB. By contrast, these experiments showed no difference in starch content between *mrc-1* and WT (Figure 11B). In fact, when comparing *mrc1* BC2 *aabb* with *mrc-1* BC2 AABB or with the WT, there was a small increase in starch content in the backcrossed mutant. The reason for the discrepancy between the results from experiments 1 – 3 compared to experiments 4 and 5 is unknown, but it may indicate that total starch content in wheat leaves is particularly variable and dependent on many environmental and physiological factors.

**Figure 11:**
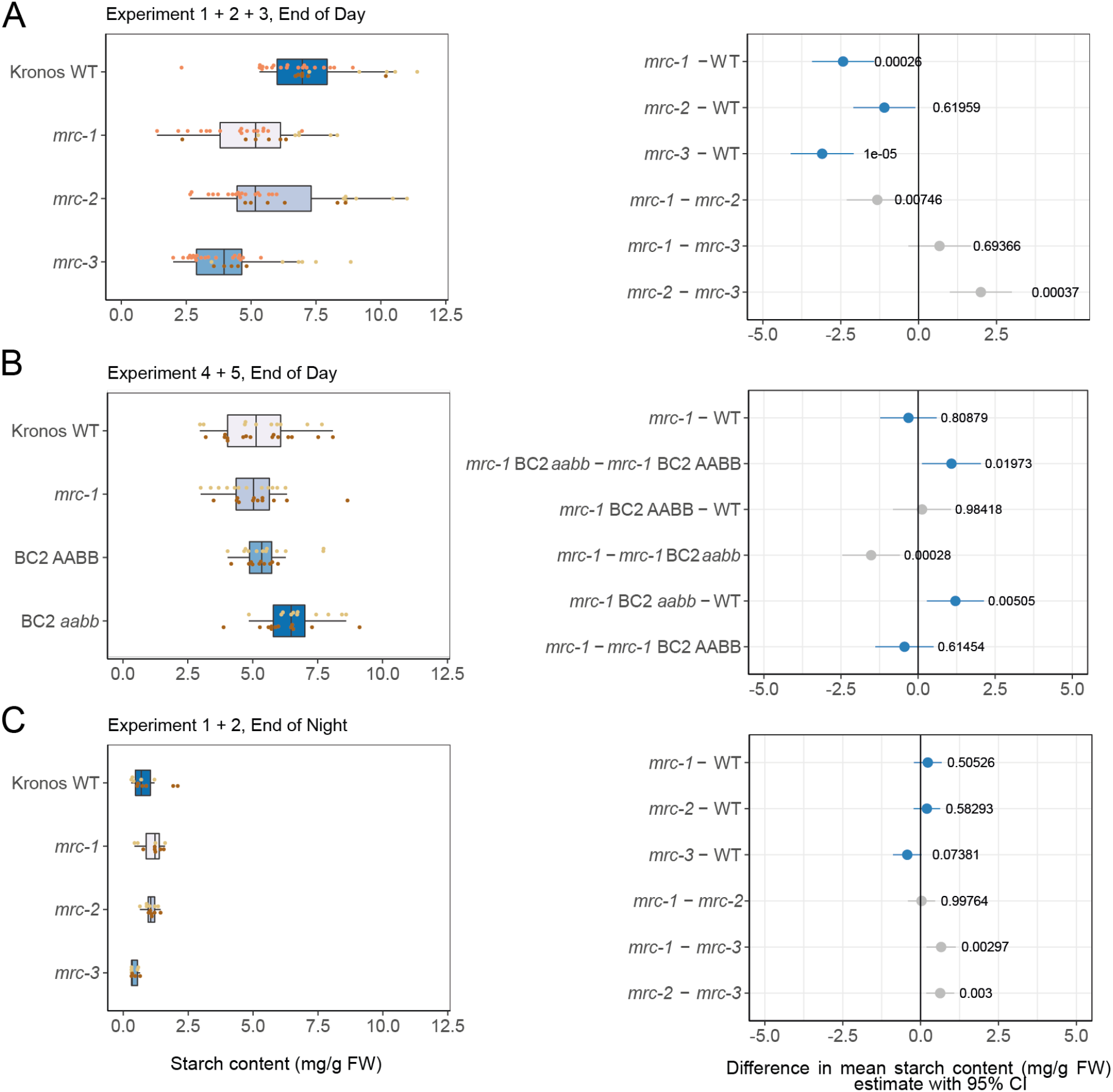
Loss of MRC has variable effects on the total end of day (ED) leaf starch content in wheat leaves. **A)** Pooled starch content data of wheat leaves harvested at the end of day (ED) in experiments 1 (dark brown, *n* = 5 – 6 per genotype), 2 (light brown, *n* = 5 – 6 per genotype) and 3 (orange, *n* = 16 – 19 per genotype). **B)** Pooled starch content data from wheat leaves in experiments 4 (dark brown, *n* = 10 – 15 per genotype) and 5 (light brown, *n* = 10 – 12 per genotype) harvested at ED. **C)** Pooled starch content data of wheat leaves harvested at the end of night (EN) in experiments 1 (dark brown, *n* = 4 – 6 per genotype) and 2 (light brown *n* = 4 – 6 per genotype) **For all panels**, raw data and boxplots are shown on the left, where dots indicate values from individual wheat plants, with colours indicating each experiment. Each box encloses the middle 50% of the distribution, the middle line is the median and the whiskers are the minimum and maximum values within 1.5 of the interquartile range. All statistical analyses were performed using a linear model with genotype and experiment as fixed effects for each panel, ANOVA and Tukey post-hoc tests. Panels on the right indicate differences in adjusted means of total starch content based on these models and pairwise comparisons of the genotypes. The difference in means is indicated by a dot, with whiskers showing the 95% confidence interval (CI) of this difference, with the corresponding p-value. For A and C, blue indicates comparisons between mutant and wild type, while grey indicates comparisons between mutants. For B, grey indicates the WT or *mrc-1* mutant compared to the backcrossed line with the equivalent genotype at the *MRC* loci, and blue indicates all other pairwise comparisons.

Most interestingly, lower starch content in *mrc* mutants compared to the WT was observed in the same batch of plants where we did not observe a significant difference in granule size between *mrc* mutants (Experiment 2, Figure 10E), and there was identical (or greater) starch content observed for *mrc-1* mutants in the same batch of plants where we observed larger granules for these mutants (Experiment 5, Figure 10F). Since the reduction in granule number described above was consistently observed in all experiments, it is plausible that the effect on granule size depends on starch content, such that the *mrc-1* mutant only produces larger granules under conditions where total starch content is equal. Pooled data from plants harvested at the end of the night (quantified in experiments 1 and 2) showed very low starch content in all three *mrc* mutants, indicating substantial nocturnal starch turnover in all genotypes, and no differences between WT and mutants (Figure 11C).

## Discussion

### A novel role for MRC in repressing B-type granule initiation during endosperm starch synthesis

Starch granule initiation remains the most enigmatic part of the starch synthesis process, where we understand little about how the diverse numbers and morphologies of starch granules are determined in our most important crops (Seung and Smith, 2019; Abt and Zeeman, 2020; Chen et al., 2021). Here, we shed new light on the temporal regulation of granule initiation in wheat endosperm, by demonstrating the unique role of the protein MRC in the repression of B-type granule initiation during early grain development.

The expression pattern of MRC during grain development is consistent with the change in onset of B-type granule initiation observed in the mutant. MRC is expressed in the endosperm between 6–10 dpa but decreases rapidly in expression between 10–15 dpa, remaining low after 15 dpa (Figure 2). This decline in expression coincides with when B-type granules start to form in the WT (between 14 –20 dpa; Figures 7 - 9). In the *mrc-1* mutant, B-type granules were initiated earlier than in the WT, already at 10 dpa (Figures 8 and 9), which could be due to the loss of B-type granule repression by MRC in early grain development. There was an increased number of starch granules in the *mrc-1* mutant compared to WT from 14 to 30 dpa (Figure 7), and this increase remained apparent in the mature grains of the backcrossed *mrc-1* mutant (Figure 4).

We propose that the early initiation of B-type granules results in drastically altered starch granule size distributions in the endosperm. The early appearance of B-type granules in the *mrc-1* mutant could present competition with A-type granules for substrates for granule growth (i.e., ADP-glucose) from an earlier stage of grain development than in the WT, resulting in a higher volume of B-type granules and lower volume of A-type granules in the mature mutant grains (Figure 5). At 8 dpa, before the appearance of B-type granules, the size distribution curves of A-type granules were almost identical between mutant and WT (Figure 8), and it was only at the later stages of grain development following B-type granule initiation that the A-type granules became smaller in the mutant compared to WT. The lack of difference in A-type granule number and size at 8 dpa suggests that the *mrc-1* mutation did not affect A-type granule initiation.

Our data show that the early B-type granule initiation alone could result in the increased proportion of B-type granule volume in mature grains, but it is difficult to make conclusions about the effect of MRC on B-type granule number specifically. The higher B-type granule volume could be caused by a combination of smaller A-type granules and larger and perhaps relatively more numerous B-type granules (Figure 12). We measured a decrease in A-type granule size and an increase in B-type granule size in our *mrc-1* and *mrc-2* mutants (Figure 5), and in some lines there was also a slight increase in total granule number (Figure 4B). However, it should be noted that there is currently no method for specifically quantifying the number of B-type granules, considering our definition of B-type granules comes from curve fitting volume/size distribution graphs from the Coulter counter, and small differences in B-type granule number may be difficult to detect as the B-type granules already make up a much larger proportion of the total granule number in WT. For *mrc-3*, a higher B-type granule volume percentage was observed (Figure 5B) without a measurable increase in B-type granule size or in total granule number (Figure 4B), but it is still possible that there is a relative increase in B-type compared to A-type granule number.

**Figure 12:**
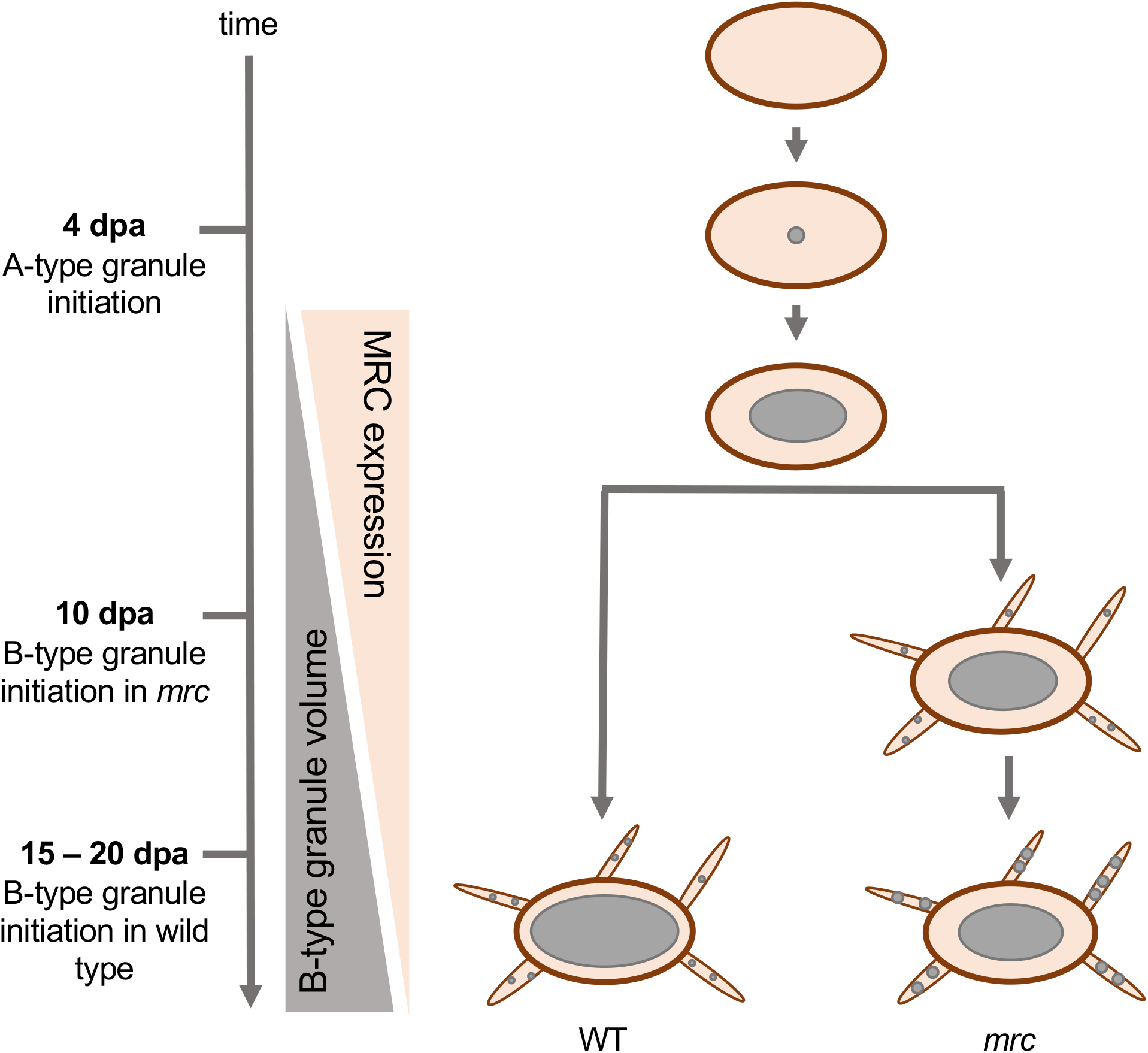
Model of MRC function in wheat developing endosperm. MRC is required for the control of the timing of B-type granule initiation during early grain development. In wild type, A-type granules initiate around 4 days post anthesis (dpa), and B-type granules initiate around 15 – 20 dpa. We propose that the expression of MRC during early endosperm development prevents the B-type granule formation, and B-type granule volume increases as MRC expression decreases after 10 dpa. This process is disrupted in mutants lacking a functional MRC protein, and granules initiate already from 10 dpa. The early initiation of B-type granules provides them with more time and substrate to grow, leading to higher volume of B-type granules in the mutant at grain maturity compared to the wild type, and a concomitant decrease in A-type granule size in the mutant.

Despite all three *mrc* mutants having an increased proportion of B-type granule volume, they differed in severity (Figure 5). The strongest effect on B-type granule percentage was seen in *mrc-1*, and since the phenotypes of the *mrc-1* backcrossed and non-backcrossed lines were similar, the severity of this line is not due to background mutations. The *mrc-1* line may have the strongest phenotype because the premature stop codon occurs earlier in the coding sequence than in *mrc-2*, and it is possible that the truncated protein in *mrc-2* is partially functional. The *mrc-3* mutant had the weakest phenotypes, suggesting the Leu289Phe mutation might also produce a partially functional protein.

The *mrc-1* wild-type segregant likely has some background mutations, as several of its phenotypes differed from the WT. The presence of background mutations is not unusual in these EMS-mutagenised wheat TILLING lines (Uauy et al., 2017), and importantly, the effects observed in the wild-type segregant were minor compared to the effects observed in the mutants.

### MRC has tissue-specific roles in promoting or repressing granule initiation

Since wheat produces starch in both leaves and endosperm, the effects of *mrc* mutations could be explored in both organs. We discovered that MRC has contrasting functions in the endosperm compared to the leaves. Rather than repressing granule initiation, MRC promotes granule initiation in the leaves, as in Arabidopsis leaves, although the mutant phenotype in wheat appears to be less severe than in Arabidopsis. In Arabidopsis *mrc* mutants, there is one large granule per chloroplast rather than the multiple smaller granules in wild-type chloroplasts (Seung et al., 2018; Vandromme et al., 2019). Wheat *mrc* chloroplasts have fewer granules per chloroplast, but there are still multiple granules in each chloroplast (Figure 10). This difference in phenotype severity is similar to those observed for the orthologs of other granule initiation proteins of Arabidopsis (BGC1 and SS4) in wheat (Hawkins et al., 2021; Watson-Lazowski et al., 2022). The *mrc-1* and *mrc-2* mutants both have fewer granules per chloroplast compared to WT, with *mrc-1* having the strongest effect, consistent with the endosperm phenotype.

The primary effect of the *mrc* mutations in leaves was the reduction in granule number per chloroplast, but we also saw varying effects on granule size. In one set of experiments, we detected larger granules in *mrc-1* compared to WT, although in another set we did not (Figure 10). This appears to be linked to whether a change in total leaf starch content was observed or not, since in both experiments, there was a reduced number of starch granules in the mutants. In the experiment where there was no difference in total starch at the end of the day, the individual (fewer) granules were larger in the mutant, as would be expected. However, in the experiment where the total starch content was lower in the *mrc-1* mutant compared to WT, the increase in granule size was not observed.

In Arabidopsis, lack of MRC does not reduce the total end of day leaf starch content; in fact, this is slightly higher in the mutant compared to the WT (Seung et al., 2018). Overall, in wheat leaves the effect of *mrc* mutations on the total end of day starch content was variable in our experiments (Figure 11). This could reflect differences in the regulation of starch turnover in wheat vs. Arabidopsis leaves, since soluble sugars make up the majority of the wheat leaf carbohydrate content and are thought to be more important as a storage reserve than starch (Nie et al., 1995; Müller et al., 2018; Watson-Lazowski et al., 2022). These differences may also contribute to the less severe effect on granule number in wheat *mrc* mutants compared to Arabidopsis leaves. The effect of MRC on leaf starch granule initiation is likely to differ depending on the developmental stage, so future studies should focus on the biochemistry of starch granule initiation across leaf and plant development to reveal a complete picture of the role of MRC and other granule initiation proteins in the wheat chloroplast. In Arabidopsis, reduced *At*SS4 abundance has a much stronger effect on granule formation in younger leaves compared to mature leaves (Crumpton-Taylor et al., 2013).

### Exploring the biochemical basis of the contrasting roles of MRC between organs

Since MRC is a long coiled-coil protein with no known enzymatic domains, it is possible that it can exert opposite effects on granule initiation by interacting with different interaction partners. The Arabidopsis *At*MRC can interact directly with *At*SS4 in a yeast two hybrid experiment (Vandromme et al., 2019), and in Arabidopsis leaves *At*MRC was also pulled down in association with *At*PTST2, the ortholog of wheat BGC1 (Seung et al., 2018). Therefore, it is possible that in wheat, protein-protein interactions also play an important role in the function of MRC, although the nature of the interactions may be quite different. Recent work has begun to uncover the distinct roles of Arabidopsis starch granule initiation orthologs in wheat, and as for MRC, they are all quite different from their role in Arabidopsis. In the endosperm, SS4 restricts granule initiation during early grain development to ensure proper A-type granule formation and mutants defective in SS4 form compound granules (Hawkins et al., 2021). Similar, but less severe, is the *bgc1* mutant in wheat which also forms irregular compound-like granules but has a higher proportion of normal-looking A-type granules than *ss4* mutants (Chia et al., 2020; Hawkins et al., 2021). BGC1 not only represses granule initiation during A-type granule formation, but also promotes B-type granule formation, as mutants with reduced gene dosage of BGC1 have fewer B-type granules than WT, with no apparent impact on A-type granules (Chia et al., 2017; Chia et al., 2020).

However, a B-type granule suppressing mechanism during early grain development, like we report here for MRC, has not previously been described, and suggests that MRC and BGC1 might have opposing roles in the wheat endosperm, despite their similar initiation promoting roles seen in leaves of Arabidopsis (Seung et al., 2017; Seung et al., 2018) and wheat (Watson-Lazowski et al., 2022). In support of this, our recent RNA-sequencing analysis of the developing wheat endosperm shows opposite expression patterns for MRC and BGC1 throughout grain development (Chen et al., 2022). Wheat SS4 also promotes leaf granule initiation, as mutants had mostly starchless chloroplasts in the leaves (Hawkins et al., 2021). Together with our finding for MRC, this suggests that there may be more distinct differences for granule initiation proteins between different organs of the same species in comparison to the same organs from different species. The Arabidopsis granule initiation proteins localise to distinct puncta in the chloroplast, but we do not yet know anything about the localisation of these wheat orthologs in the chloroplast or amyloplast. In the future, exploring the protein localisation, protein-protein interactions, and the connection between the two in both tissues could reveal what underpins these distinct functions of MRC.

The specific effect of the L289F mutation in the *mrc-3* mutant on endosperm starch may provide an important clue in these studies, since *mrc-3* had increased B-type granule percentage in the endosperm, but no differences in granule number in leaves. Determining whether the L289F mutation affects protein conformation, interactions and localisation might help dissect the differences in MRC function in leaves versus endosperm.

### MRC as a gene target for biotechnological modification of starch granule size

There is significant industrial interest in manipulating starch granule size in crop species, as granule size affects the physico-chemical properties of starch and digestibility (Jobling, 2004; Lindeboom et al., 2004; Chen et al., 2021; Li et al., 2021). Our results establish MRC as a promising gene target for modifying starch granule size distribution in wheat, specifically to achieve smaller starch granules and a narrower granule size distribution range than conventional cultivars. Small granules are more efficiently digested *in vitro* than large granules, due to their larger surface area to volume ratio (Dhital et al., 2010). Possible uses for wheat *mrc* starch within the food industry include pasta making, where more B-type granules positively affect pasta quality due to their higher rate of water absorption (Soh et al., 2006). Wheat *mrc* starch may also be useful in industrial applications like papermaking and biodegradable plastics, where small granules are desirable (Lindeboom et al., 2004). Functional tests can be directly performed on our material to provide proof of concept that the altered granule size distribution in the *mrc* mutants improves grain/starch quality.

MRC is a promising target because different granule size can be achieved without accompanying effects on overall plant growth (Figure 3), on grain weight and total starch content (Figure 4), on starch granule shape (Figure 6), or on amylopectin structure and amylose content (Supplemental Figure 3). It is also ideal that the B-genome homeolog has become a pseudogene (Figure 1), meaning that only one or two homeologs need to be mutated in durum and bread wheat respectively to achieve an effect on granule size. Further, we have also demonstrated that different mutations in MRC can be used to fine-tune the volume of B-type granules in endosperm starch, such as those in *mrc-2* and *mrc-3* to achieve moderate increases, and *mrc-1* to achieve larger increases. Along this line, we are currently investigating whether overexpression of MRC can be used to reduce the volume of B-type granules.

Since the repression of B-type granule initiation is likely to be a role specific to Triticeae species carrying a bimodal size distribution of endosperm starch, it remains to be determined what the role of MRC is in cereal species that do not have a bimodal distribution of starch granules, such as those that have compound granules (e.g. in rice). Also, oats have a bimodal distribution of starch granules, with large compound granules and smaller simple granules. However, in oat, the smaller granules initiate at the same time as the larger compound granules (Saccomanno et al., 2017), and it would therefore be interesting to determine if differences in MRC function play a role in timing the initiation of the small granules during oat endosperm development. Exploring the role of MRC in multiple crop species would therefore not only reveal the molecular differences that result in distinct spatiotemporal patterns of granule initiation between species, but also potentially increase its biotechnological potential.

## Materials and Methods

### TaMRC bioinformatic analysis

To characterize *TtMRC-B1* in tetraploid wheat, we aligned the whole genome-sequencing reads of *Triticum dicoccoides* (n = 10) and *Triticum turgidum ssp. durum* (n = 10) from Zhou et al., 2020 against the ‘tetraploid’ version of the Ref-Seqv1.0 Chinese Spring assembly (Appels et al., 2018). We used HiSat2-v-2.1.0 (Kim et al., 2019) with the default settings and visualized the read alignments on the genetic signatures of retrotransposon insertion on *MRC-B1* (Figure 1B) using Integrated Genomics Viewer (Robinson et al., 2011). Phylogenetic analyses were performed as described in Seung et al., (2018).

### Plant materials and growth

EMS mutants of tetraploid wheat (*Triticum turgidum* cv. Kronos) carrying mutations in *TtMRC-A1* and the chromosome 6B pseudogene were identified from the wheat *in silico* TILLING database (http://www.wheat-tilling.com; Krasileva et al., 2017) and obtained from the John Innes Centre Germplasm Resource Unit. The selected mutants for *TtMRC-A1* were Kronos3272(K3272), Kronos598(K598) and Kronos4681(K4681); while Kronos4305(K4305) and Kronos3078(K3078) were selected for the 6B pseudogene. From these mutants, we generated three different sets of lines. The *mrc-1* lines descend from a cross between K3272 and K3078, while the *mrc-2* lines descend from a cross between K4681 and K4305. For both crosses, *aa*BB, AA*bb* and *aabb* genotypes were obtained in the F2 generation. The *mrc-3* lines are the original K598 mutants. The KASP markers used to genotype the mutations are provided in Supplemental Table 3.

For experiments on grains and leaves, plants were grown in soil in a controlled environment room with fluorescent lamps supplemented with LED panels. The chambers were set to provide a 16-h light at 300 μmol photons m^-2^ s^-1^ and 20°C, and 8-h dark at 16°C, with relative humidity of 60%. Grains from the first three tillers were harvested from mature, dry spikes (approximately 4 months after sowing). Leaves were harvested 10 days after germination, when two leaves were present. The two leaves were pooled for starch quantification, and a section from the middle of the older leaf was used for light microscopy.

### Grain morphometrics

The number of grains harvested per plant, plus grain area and thousand grain weight were quantified using the MARViN seed analyser (Marvitech GmbH, Wittenburg). Multiple grains from each plant (15 - 88 individual grains per plant) were measured for grain area to calculate the average value for each plant, and these values were used in the plots of Figure 3F and the analysis of Supplemental Table 2D.

### Starch purification from mature grains or developing endosperm

Starch was purified from mature grains using 3-6 grains per extraction. Dry grains were soaked overnight at 4°C in 5 mL of sterile water. The softened grains were homogenised in 10 mL sterile water using a mortar and pestle, and the homogenate was filtered through a 100 μm mesh. The starch was pelleted by centrifugation at 3,000*g* for 5 minutes, and resuspended in 2 mL of water. The resuspended starch was loaded on top of a 5 mL 90% Percoll (Sigma) cushion buffered with 50 mM Tris-HCl, pH 8, and was spun at 2,500*g* for 15 minutes. We verified that no intact granules were left in the Percoll interface after the spin. The starch pellet was washed twice with wash buffer (50 mM Tris-HCl, pH 6.8; 10 mM EDTA; 4% SDS; and 10 mM DTT), then three times with water, followed by a final wash in absolute ethanol. The starch was then air dried overnight.

For starch extraction from developing endosperm, developing grains were harvested at the indicated timepoints and were snap frozen in liquid nitrogen and stored at −80°C until analysis. Each grain was thawed just prior to extraction and the endosperm was carefully dissected and placed into a chilled tube and weighed. The tissue was then homogenised in sterile water with a pestle, then filtered through a 60 μm mesh. The pellet was washed three times in 90% Percoll (Sigma) buffered with 50 mM Tris-HCl, pH 8, then three times with wash buffer (as above), followed by three times with water.

### Coulter counter analysis of starch granule size and number

For profiles of granule size distribution, purified starch was suspended in Isoton II diluent (Beckman Coulter) and analysed with a Multisizer 4e Coulter counter fitted with a 70 μm aperture (Beckman Coulter). Granules were counted either in volumetric mode, measuring 1 mL from a total 100 mL volume preparation containing 20 μL of purified starch or set to count at least 100,000 granules. For calculations of granule counts in volumetric mode, the number of granules per mg grain weight was back calculated to the total starting grain weight. The granules were sized, with the Coulter counter collecting the data using logarithmic bins for the granule diameter (standard settings).

For each plant, to calculate the mean A- and B-type granule size, as well as relative B-type granule volume, we fitted a mixed bimodal gaussian curve to the distribution using R (https://github.com/JIC-CSB/coulter_counter_fitting). As the data collection on the Coulter counter is set to logarithmic bins on the x-axis, for these calculations and for the traces in Figures 5A, 8A, and Supplemental Figure 3, we transformed the x-axis to even bins, by changing the y-axis to volume percentage density (volume percentage for each bin divided by bin width). For each of the extracted phenotypes of mean A- and B-type granule size and relative B-type granule volume, we fitted individual linear models and performed a one-way ANOVA and Tukey post-hoc tests for pairwise comparisons of the genotypes, using the lm() and emmeans() functions in R, from the ‘stats’ and ‘emmeans’ packages. Nine individual plants per genotype were used for this experiment.

### Light and electron microscopy

For light microscopy of starch granules in leaf chloroplasts, leaf material (1.5 mm x 1.5 mm squares) was harvested and fixed in 2.5% glutaraldehyde in 0.05 M sodium cacodylate, pH 7.4, which was vacuum infiltrated into the leaf. The segments were post-fixed with osmium, then dehydrated in an ascending ethanol series, and embedded in LR white resin using EM TP embedding machine (Leica). Semi-thin sections (0.5 μm thick) were produced from the embedded leaves using a glass knife and were dried onto PTFE-coated slides. Chloroplasts and cell walls were stained using toluidine blue stain (0.5% toluidine blue ‘O’, 0.5% sodium borate) for 1 min. Starch was stained using reagents from the Periodic Acid-Schiff staining kit (Abcam): using a 30 min incubation with periodic acid solution, followed by 5 min with Schiff’s solution. Then, chloroplasts and cell walls were again stained using toluidine blue stain for 1 min. The sections were mounted with Histomount (National Diagnostics) and imaged on a DM6000 microscope with 63X oil immersion lens (Leica).

For light microscopy of endosperm sections from mature grains, thin sections (1 μm thick) of mature grains were made using a microtome fitted with a glass knife. Sections were mounted onto a glass slide and stained with 3% Lugol’s iodine solution (Sigma) prior to imaging on a DM6000 (Leica) or AxioObserver Z1 (Zeiss) microscope.

For light/electron microscopy of developing endosperm tissue, developing grains (15 dpa) were harvested into 4% paraformaldehyde, 2.5% glutaraldehyde in 0.05 M sodium cacodylate, pH 7.4. The osmium post-fixation, dehydration and embedding into LR white resin was done as described above for leaves. For light microscopy, semi-thin sections were stained with toluidine blue and imaged as described above.

For transmission electron microscopy (TEM), ultra-thin sections (approximately 90 nm) were produced from the embedded grains by sectioning with a diamond knife using a Leica UC7 ultramicrotome (Leica, Milton Keynes). The sections were picked up on 200 mesh copper grids which had been formvar and carbon coated, then stained with 2% (w/v) uranyl acetate for 1hr and 1% (w/v) lead citrate for 1 minute, washed in distilled water and air dried. The grids were viewed in a FEI Talos 200C transmission electron microscope (FEI UK Ltd, Cambridge, UK) at 200kV and imaged using a Gatan OneView 4K x 4K digital camera (Gatan, Cambridge, UK) to record DM4 files.

For scanning electron microscopy (SEM): For imaging starch granules, a drop of purified starch suspended in water (5 mg/mL) was air-dried onto a glass coverslip attached onto an SEM stub. For imaging sections through developing endosperm, harvested grains were fixed in 2.5% glutaraldehyde in 0.05 M sodium cacodylate, pH 7.4. The fixative was removed by washing with 0.05 M sodium cacodylate, pH 7.4, after which the grains were dehydrated in an ascending ethanol series, and then subjected to critical point drying in a CPD300 instrument (Leica) according to the manufacturer’s instructions. Thick transverse sections were produced from the dried grains and were glued onto SEM stubs. All stubs were sputter coated with gold and observed using either a Supra 55 VPFEG (Zeiss) or Nova NanoSEM 450 (FEI) SEM instrument.

### Analysis of leaf light microscopy images

For quantification of the number of granules per chloroplast, light microscopy images from three individual plants of each of experiments 2 and 5 were used (Supplemental Table 4). Using the cell counter plug-in in ImageJ, the number of granules per chloroplast were counted manually, until 200 – 250 chloroplasts were reached. Cells were chosen across the section, where most of the granules were stained well and clearly visible, and cells directly next to vasculature were avoided. It should be noted that these are counts of granule sections, rather than the actual numbers of granules per chloroplast, as the actual number cannot be determined from two-dimensional sections. As much as possible, all chloroplasts in a cell were counted to avoid bias, although occasionally a few chloroplasts where the granules were unclear had to be omitted. Both lighter and darker stained granules were counted, but occasionally light-staining patches that were difficult to discern as granules were observed in all samples, and these were not included in the count.

For quantification of the granule size, we analysed the same samples as for the granules per chloroplast, using the ‘analyse particles’ function in ImageJ, and manually adjusted the thresholding in an image until the dark granules in the images were correctly separated. This was done for several images across the section, until 100 – 400 granules were reached for each plant. As much as possible, we removed instances where the thresholding had fused multiple granules in proximity by checking obvious outliers against the original images.

For the statistics of the quantification of the number of granules per chloroplast, we consulted the Statistical Services Centre Ltd (Reading, UK), who wrote the statistical analysis script in R for us, available at [https://github.com/Jiaawen/2022_MRC_wheat_Rscripts/tree/main/Fig10]. We used a negative binomial mixed effects model, since we were analysing count data and wanted to account for both the random effect of biological replicate and the frequency distribution of number of granules per chloroplast. We used the mixed_model() function with ‘family=negative.binomial’ from the ‘GLMMadaptive’ package, and the emmeans() function from the ‘emmeans’ package for pairwise comparisons in R. Our data had overdispersion when fitting a Poisson model but not when using a negative binomial. We used individual plants as the random effect (for both experiments: 3 plants for each of 4 genotypes – 12 plants total), and genotype as fixed effect.

For the statistics of the granule area, we used a linear mixed effects model, also with the individual plants as random effect and genotype as fixed effect. We used the lmer() function from the ‘lmerTest’ package, and the emmeans() function from the ‘emmeans’ package for pairwise comparisons.

### Quantification of starch content in leaves and endosperm

Starch was quantified in leaf tissue as previously described (Smith and Zeeman, 2006). Briefly, frozen leaf tissue was ground into a powder with a ball mill and then extracted with perchloric acid. Starch in the insoluble fraction of the extraction was gelatinised at 95°C and digested to glucose with α-amylase (Megazyme) and amyloglucosidase (Roche). The glucose released was measured using the hexokinase/glucose-6-phosphate dehydrogenase method (Roche). Starch content (in glucose equivalents) was calculated relative to the original dry weight of the analysed grains.

Five individual experiments were done to quantify the total end of day leaf starch content (Supplemental Table 4). One set of genotypes was measured in experiments 1 – 3: WT, *mrc-1, mrc-2, mrc-3*. Another set was measured in experiments 4 and 5: WT, *mrc-1*, the double backcrossed *mrc-1* BC2 *aabb* and the wild-type segregant from that backcross, *mrc-1* BC2 AABB. Therefore, in the statistical analysis experiments 1, 2, 3 were pooled together and experiments 4 and 5 were pooled together. An ANOVA using a fixed effects model showed no interaction between the experiment effect and genotype effect, and experiment and genotype were set as fixed effects in our linear model. We fitted individual linear models for end of day starch in experiments 1 – 3, end of day starch in experiments 4 & 5 and end of night starch in experiments 1 & 2. Then we performed ANOVA and Tukey post-hoc tests using these linear models for pairwise comparisons of the genotypes. For the details of number of replicates, see Supplemental Table 4. We used the lm() function from the ‘stats’ package in R for the linear models, and used the emmeans() function from the ‘emmeans’ package for calculating adjusted means and pairwise comparisons.

A similar method to leaves was used for starch quantification in grains. Mature grains (5-6 grains) were soaked overnight at 4°C in 5 mL of sterile water and were homogenised using a mortar and pestle. Developing endosperm tissue was extracted in 1 mL of sterile water with the pestle. Insoluble material in an aliquot of the homogenate was collected by centrifugation at 5,000*g* for 5 mins, then washed once in 0.7 M perchloric acid, once in sterile water, then three times in 80% ethanol. The pellet was then resuspended in water. Starch in the pellet was gelatinised by heating at 95°C for 15 min, then digested using α-amylase (Megazyme) and amyloglucosidase (Roche).

### Analysis of amylopectin structure and amylose content

Amylopectin structure and amylose content were analysed using purified starch. Amylopectin structure in terms of chain length distribution was quantified using High Performance Anion Exchange Chromatography with Pulsed Amperometric Detection (HPAEC-PAD) (Blennow et al., 1998). For amylose content, granules were dispersed in DMSO and quantified using an iodine-binding method (Warren et al., 2016).

### Statistics

All statistical analyses were done in R version 4. Overall, we used the emmeans() function from the ‘emmeans’ package throughout for pairwise comparisons. For most experiments, we used linear models using the lm() function from the ‘stats’ package and did one-way ANOVAs with Tukey post-hoc tests. Details for individual experiments where we used other models with more than a single fixed effect are described above for the relevant sections.

Tables 1, 2 and Supplemental Table 2 also describe details of statistics for individual experiments.

## Accession numbers

The accession numbers corresponding to the genes investigated in this study are*: Tt*MRC-A1 (TRITD6Av1G081580), *Ta*MRC-A1 (TraesCS6A02G180500), *Ta*MRC-D1 (TraesCS6D02G164600).

## Acknowledgements

The authors thank the John Innes Centre (JIC) Horticultural Services for providing growth facilities and maintenance of plant material, and JIC Bioimaging for providing access to microscopes. We thank Prof. Alison Smith for critically reading this manuscript. This work was funded through a John Innes Foundation (JIF) Chris J. Leaver Fellowship (to D.S), a Biotechnology and Biological Sciences Research Council (BBSRC, UK) Future Leader Fellowship BB/P010814/1 (to D.S.), a JIF Rotation Ph.D. studentship (to J.C. and Y.C.) and BBSRC Institute Strategic Programme grants BBS/E/J/000PR9790, BBS/E/J/000PR9799 and BB/P016855/1 (to the John Innes Centre).

## Author contributions

J.C. and D.S. conceived the study. J.C., Y.C., A.W-L., F.J.W., A.B., C.U., and D.S. designed the research. J.C., Y.C., A.W-L., E.H., J.E.B., B.F., R.D.B., K.C., F.J.W., A.B., C.U., and D.S. performed the research and analysed data; J.C. and D.S. wrote the article with input from all authors.

## Supplemental data

Supplemental File 1. *Aegilops speltoides* MRC gene model

Supplemental Figure 1. Phylogenetic analysis of wheat MRC sequences.

Supplemental Figure 2. Chain length distribution and amylose content of the *mrc* mutants.

Supplemental Figure 3. The 6B pseudogene does not contribute to granule size distribution in wheat endosperm starch.

Supplemental Table 1. Reads mapped to retrotransposon insertion genetic signatures on *MRC-B1*.

Supplemental Table 2. Pairwise comparisons of wheat growth phenotypes.

Supplemental Table 3. KASP markers for genotyping the wheat mutants.

Supplemental Table 4. Summary of leaf starch quantification experiments.

