## Supplemental Figures for "The plastidial protein MRC promotes starch granule initiation in wheat leaves but delays B-type granule initiation in the endosperm"

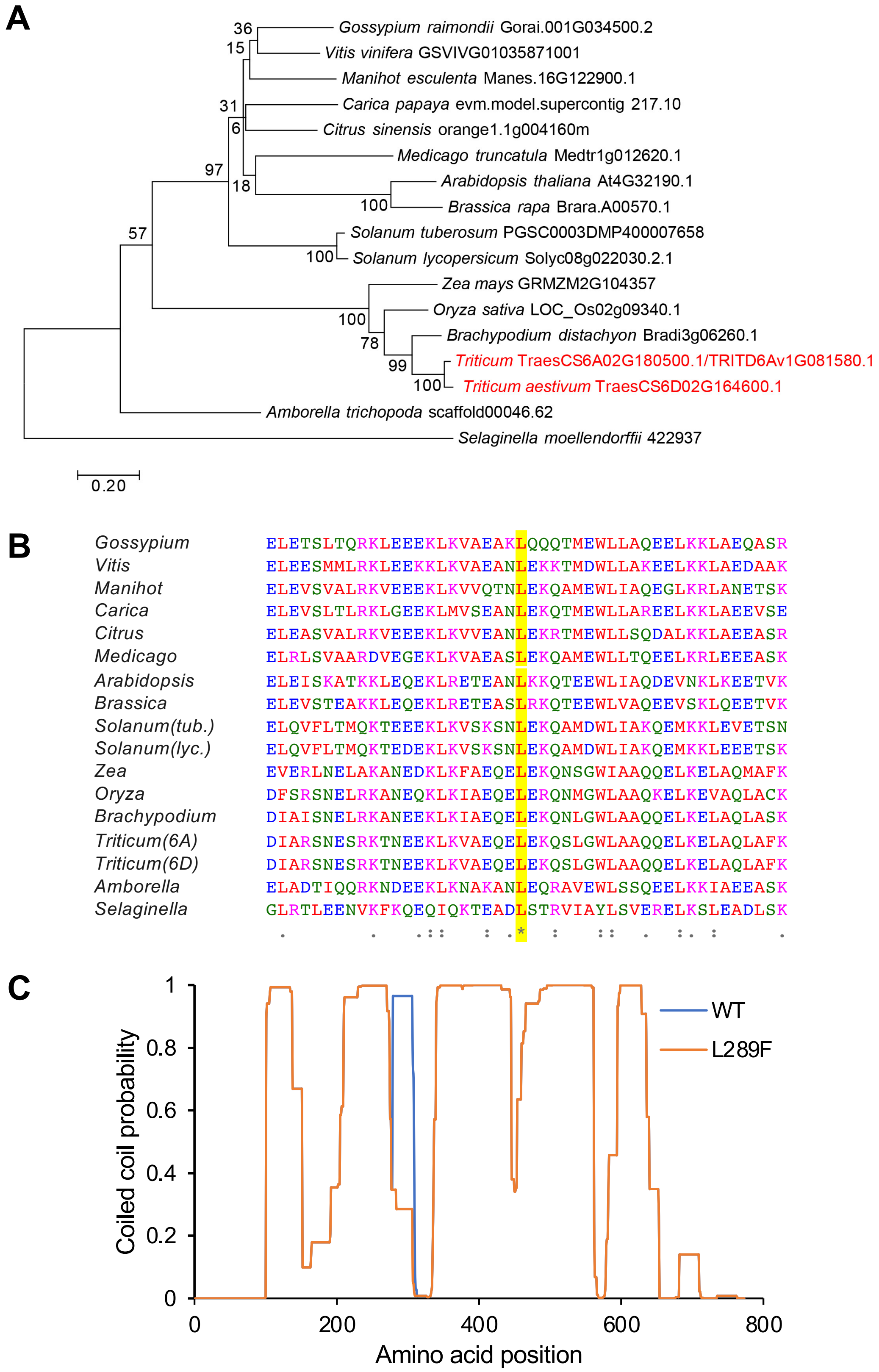


**Supplemental Figure 1. Phylogenetic analysis of wheat MRC sequences. A)** Phylogenetic tree of MRC orthologs. The amino acid sequences of the orthologs identified in Seung *et al.* 2018 were aligned with the wheat sequences. A maximum likelihood tree was assembled with 1000 bootstraps. Bootstrap values are shown next to each node. Branch lengths indicate the number of substitutions per site. **B)** Multiple sequence alignment of MRC orthologs generated with Clustal O. The region surrounding Leu289 (highlighted in yellow) is shown. Symbols under the alignment indicate conserved residues with: complete identity (*), highly similar properties (:) or weakly similar properties (.). Colours of residues represent side chain properties: hydrophobic (red), acidic (blue), basic (magenta) and other (green). **C)** Coiled coil prediction of *Ta*MRC-A1 wild type (WT) sequence and the Leu289Phe (L289F) mutant using COILS/PCOILS. The probability was calculated using the 28 amino acid window.

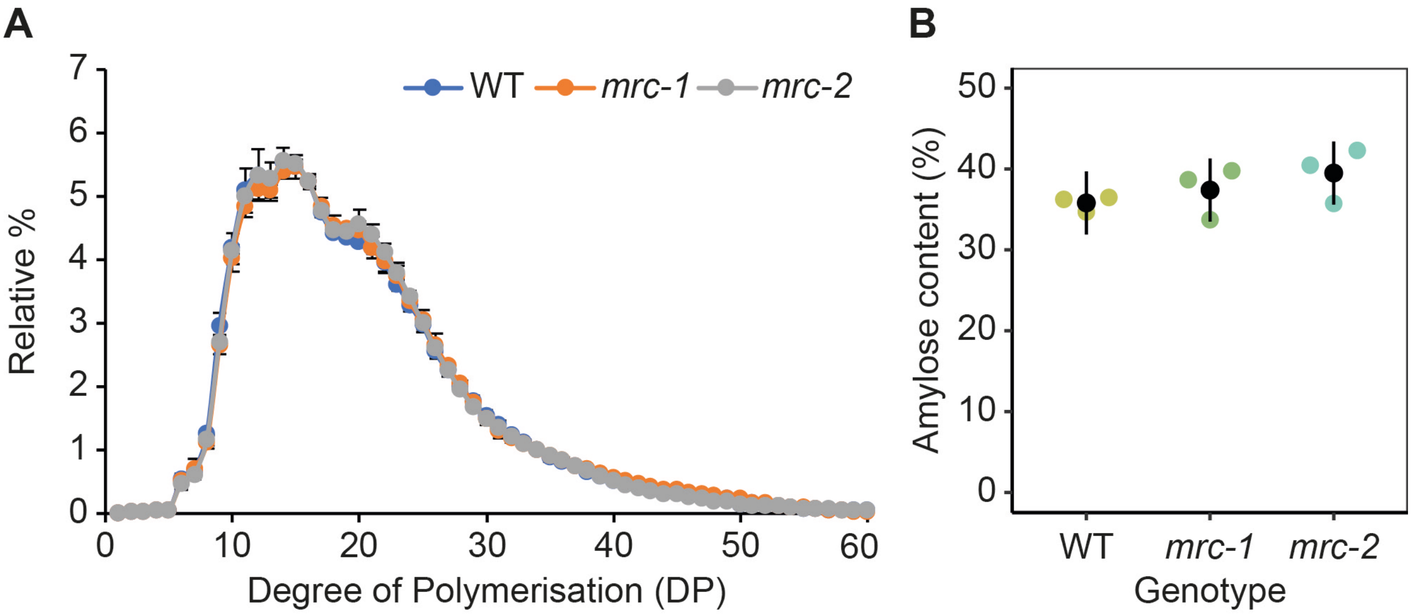


**Supplemental Figure 2. Chain length distribution and amylose content of the *mrc* mutants.**
**A)** Chain length distribution of *mrc-1* and *mrc-2* starch. Purified starch was debranched and analysed with High Performance Anion Exchange Chromatography with Pulsed Amperometric Detection (HPAEC-PAD). The area of peaks corresponding to chains of each degree of polymerisation (DP) was expressed as a percentage of the summed peak area for DP 1–60. Values are the mean ± SEM from three replicate measurements. **B)** Amylose content of of *mrc-1* and *mrc-2* starch quantified using iodine colourimetry. Values represent mean ± 95% CI from three replicate measurements.

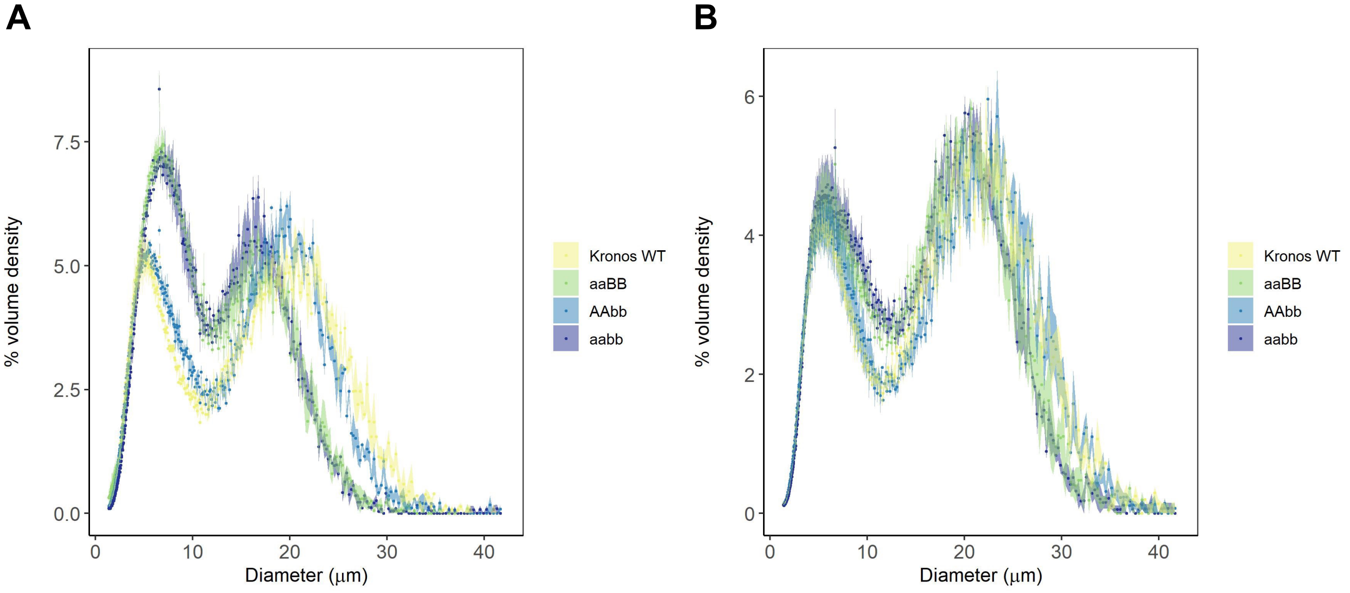


**Supplemental Figure 3. The 6B pseudogene does not contribute to granule size distribution in wheat endosperm starch.** Size distributions were determined by measuring at least 100,000 granules per replicate with a Coulter counter, and are plotted here with evenly binned x-axes. Data points are mean values from 3 individual plants of each genotype (3 grains from each plant), with the standard error of the mean shown as a shaded ribbon. **A)** Starch from genotypes isolated from the *mrc-1* cross: the single A homeolog mutant (*aa*BB), the single B pseudogene mutant (AA*bb*), and the double mutant (*aabb*). **B)** Same as A), but with genotypes from the *mrc-2* cross.
