## Supplemental Tables for "The plastidial protein MRC promotes starch granule initiation in wheat leaves but delays B-type granule initiation in the endosperm"

**Supplemental Table 1. Reads mapped to genetic signatures of retrotransposon insertion in *MRC-B1.*** *Triticum turgidum ssp. durum* and *Triticum dicoccoides* species (Zhou *et al.* 2020) were aligned to the 'tetraploid' version of the Chinese Spring reference genome, and the number of reads mapped to each of the three genetic signatures in each line is indicated.

| Zhou *et al.* (2020) Line ID | Subspecies | 5' UTR and exon two junction | 5' junction of the retrotransposon insertion | 3' junction of the retrotransposon insertion |
|---|---|---|---|---|
| PI 24493 | Triticum turgidum L. ssp. durum (Desf.) Husn. | 3 | 4 | 2 |
| PI 166327 | Triticum turgidum L. ssp. durum (Desf.) Husn. | 1 | 7 | 8 |
| PI 178143 | Triticum turgidum L. ssp. durum (Desf.) Husn. | 1 | 1 | 7 |
| PI 192051 | Triticum turgidum L. ssp. durum (Desf.) Husn. | 4 | 4 | 2 |
| PI 623461 | Triticum turgidum L. ssp. durum (Desf.) Husn. | 3 | 0 | 6 |
| PI 624129 | Triticum turgidum L. ssp. durum (Desf.) Husn. | 2 | 9 | 8 |
| PI 624388 | Triticum turgidum L. ssp. durum (Desf.) Husn. | 4 | 4 | 4 |
| PI 625273 | Triticum turgidum L. ssp. durum (Desf.) Husn. | 2 | 6 | 7 |
| PI 626483 | Triticum turgidum L. ssp. durum (Desf.) Husn. | 3 | 5 | 0 |
| PI 627514 | Triticum turgidum L. ssp. durum (Desf.) Husn. | 4 | 5 | 6 |
| PI 627942 | Triticum turgidum L. ssp. durum (Desf.) Husn. | 4 | 4 | 5 |
| PI 627996 | Triticum turgidum L. ssp. durum (Desf.) Husn. | 2 | 2 | 3 |
| PI 428016 | Triticum turgidum L. ssp. dicoccoides (Korn. ex Asch. & Graebn.) Thell. | 2 | 4 | 5 |
| PI 466933 | Triticum turgidum L. ssp. dicoccoides (Korn. ex Asch. & Graebn.) Thell. | 4 | 4 | 4 |
| PI 466970 | Triticum turgidum L. ssp. dicoccoides (Korn. ex Asch. & Graebn.) Thell. | 2 | 4 | 0 |
| PI 471062 | Triticum turgidum L. ssp. dicoccoides (Korn. ex Asch. & Graebn.) Thell. | 5 | 3 | 7 |
| PI 487254 | Triticum turgidum L. ssp. dicoccoides (Korn. ex Asch. & Graebn.) Thell. | 3 | 2 | 0 |
| PI 428098 | Triticum turgidum L. ssp. dicoccoides (Korn. ex Asch. & Graebn.) Thell. | 4 | 1 | 1 |
| PI 428138 | Triticum turgidum L. ssp. dicoccoides (Korn. ex Asch. & Graebn.) Thell. | 5 | 8 | 2 |
| TRI 11505 | Triticum turgidum L. ssp. dicoccoides (Korn. ex Asch. & Graebn.) Thell. | 3 | 2 | 4 |
| PI 428041 | Triticum turgidum L. ssp. dicoccoides (Korn. ex Asch. & Graebn.) Thell. | 2 | 2 | 1 |
| PI 428071 | Triticum turgidum L. ssp. dicoccoides (Korn. ex Asch. & Graebn.) Thell. | 3 | 2 | 4 |

**Supplemental Table 2. Pairwise comparisons of wheat growth phenotypes. A, B)** Ratios of pairwise comparisons between genotypes, based on a Poisson regression model with post-hoc tests performed on the log scale and intervals back-transformed from the log scale. For the post-hoc tests, p-value and 95% confidence intervals were adjusted using Bonferroni method. Statistics were done in R, using the glm() function from the 'GLMMadaptive' package. **C, D)** Differences of pairwise comparisons between genotypes, based on a linear regression model and one-way ANOVA with Tukey post-hoc test. Statistics were done in R using the lm() function from the 'stats' package. All pairwise comparisons were done using the emmeans() function from the 'emmeans' package. Comparisons where p<0.05 are highlighted in grey. SE = standard error, 95% CI = 95% confidence interval, df = degrees of freedom.

A)

| **Tiller number per plant** | | | | | | |
|---|---|---|---|---|---|---|
| contrast | ratio | SE | z.ratio | p.value | - 95% CI | + 95 CI |
| WT / *mrc-1* | 0.82 | 0.152 | -1.07 | 1 | 0.48 | 1.41 |
| WT / *mrc-2* | 0.78 | 0.142 | -1.39 | 1 | 0.45 | 1.32 |
| WT / *mrc-3* | 0.91 | 0.173 | -0.48 | 1 | 0.52 | 1.59 |
| WT / *mrc-1* BC2 *AABB* | 0.91 | 0.173 | -0.48 | 1 | 0.52 | 1.59 |
| WT / *mrc-1* BC2 *aabb* | 1.01 | 0.196 | 0.06 | 1 | 0.57 | 1.79 |
| *mrc-1/ mrc-2* | 0.95 | 0.159 | -0.34 | 1 | 0.58 | 1.55 |
| *mrc-1 /mrc-3* | 1.11 | 0.195 | 0.61 | 1 | 0.67 | 1.86 |
| *mrc-1 / mrc-1* BC2 *AABB* | 1.11 | 0.195 | 0.61 | 1 | 0.67 | 1.86 |
| *mrc-1 / mrc-1* BC2 *aabb* | 1.23 | 0.222 | 1.16 | 1 | 0.73 | 2.09 |
| *mrc-2 / mrc-3* | 1.18 | 0.203 | 0.95 | 1 | 0.71 | 1.95 |
| *mrc-2 / mrc-1* BC2 *AABB* | 1.18 | 0.203 | 0.95 | 1 | 0.71 | 1.95 |
| *mrc-2 / mrc-1* BC2 *aabb* | 1.30 | 0.232 | 1.49 | 1 | 0.77 | 2.20 |
| *mrc-3 / mrc-1* BC2 *AABB* | 1 | 0.180 | 1.36E-15 | 1 | 0.59 | 1.69 |
| *mrc-3 / mrc-1* BC2 *aabb* | 1.11 | 0.204 | 0.55 | 1 | 0.64 | 1.90 |
| mrc-1 BC2 *AABB* / mrc-1 BC2 *aabb* | 1.11 | 0.204 | 0.55 | 1 | 0.64 | 1.90 |

B)

| **Grain number per plant** | | | | | | |
|---|---|---|---|---|---|---|
| contrast | ratio | SE | z.ratio | p.value | - 95% CI | + 95 CI |
| WT / *mrc-1* | 1.05 | 0.045 | 1.026 | 1 | 0.92 | 1.19 |
| WT / *mrc-2* | 0.91 | 0.038 | -2.335 | 0.293151 | 0.80 | 1.03 |
| WT / *mrc-3* | 1.15 | 0.051 | 3.117 | 0.027373 | 1.01 | 1.31 |
| WT / *mrc-1* BC2 *AABB* | 1.28 | 0.058 | 5.464 | 6.95E-07 | 1.12 | 1.46 |
| WT / *mrc-1* BC2 *aabb* | 1.04 | 0.045 | 1.005 | 1 | 0.92 | 1.18 |
| mrc-1 /*mrc-2* | 0.87 | 0.036 | -3.452 | 0.008351 | 0.77 | 0.98 |
| mrc-1 / *mrc-3* | 1.10 | 0.048 | 2.147 | 0.477097 | 0.97 | 1.24 |
| *mrc-1 / mrc-1* BC2 *AABB* | 1.23 | 0.055 | 4.554 | 7.88E-05 | 1.08 | 1.40 |
| *mrc-1 / mrc-1* BC2 *aabb* | 1.00 | 0.042 | -0.021 | 1 | 0.88 | 1.13 |
| *mrc-2 / mrc-3* | 1.26 | 0.053 | 5.583 | 3.55E-07 | 1.12 | 1.43 |
| *mrc-2 / mrc-1* BC2 *AABB* | 1.41 | 0.061 | 7.956 | 2.67E-14 | 1.24 | 1.60 |
| *mrc-2 / mrc-1* BC2 *aabb* | 1.15 | 0.047 | 3.431 | 0.009028 | 1.02 | 1.30 |
| *mrc-3 / mrc-1* BC2 *AABB* | 1.12 | 0.051 | 2.418 | 0.23417 | 0.98 | 1.28 |
| *mrc-3 / mrc-1* BC2 *aabb* | 0.91 | 0.040 | -2.168 | 0.452384 | 0.80 | 1.03 |
| mrc-1 BC2 *AABB* / mrc-1 BC2 *aabb* | 0.81 | 0.036 | -4.575 | 7.13E-05 | 0.71 | 0.93 |

C)

| Thousand Grain Weight (g) | | | | | | | |
|---|---|---|---|---|---|---|---|
| contrast | difference | SE | df | t.ratio | p.value | - 95% CI | + 95 CI |
| WT - *mrc-1* | 0.597 | 2.768 | 53 | 0.215819 | 0.999932 | -7.588 | 8.782 |
| WT - *mrc-2* | 3.031 | 2.768 | 53 | 1.094812 | 0.881287 | -5.154 | 11.216 |
| WT - *mrc-3* | -5.275 | 2.768 | 53 | -1.90551 | 0.410261 | -13.460 | 2.910 |
| WT - *mrc-1* BC2 *AABB* | -8.684 | 2.768 | 53 | -3.13658 | 0.031545 | -16.869 | -0.499 |
| WT - *mrc-1* BC2 *aabb* | 1.104 | 2.768 | 53 | 0.3986 | 0.998628 | -7.081 | 9.289 |
| *mrc-1 - mrc-2* | 2.433 | 2.695 | 53 | 0.903079 | 0.944029 | -5.533 | 10.400 |
| *mrc-1 - mrc-3* | -5.873 | 2.695 | 53 | -2.17945 | 0.264477 | -13.840 | 2.094 |
| *mrc-1 - mrc-1* BC2 *AABB* | -9.281 | 2.695 | 53 | -3.44427 | 0.013618 | -17.248 | -1.314 |
| *mrc-1 - mrc-1* BC2 *aabb* | 0.506 | 2.695 | 53 | 0.187789 | 0.999966 | -7.461 | 8.473 |
| *mrc-2 - mrc-3* | -8.306 | 2.695 | 53 | -3.08253 | 0.036314 | -16.273 | -0.340 |
| *mrc-2 - mrc-1* BC2 *AABB* | -11.715 | 2.695 | 53 | -4.34735 | 0.000851 | -19.681 | -3.75 |
| *mrc-2 - mrc-1* BC2 *aabb* | -1.927 | 2.695 | 53 | -0.71529 | 0.979257 | -9.894 | 6.039 |
| *mrc-3 - mrc-1* BC2 *AABB* | -3.408 | 2.695 | 53 | -1.26481 | 0.802435 | -11.375 | 4.558 |
| *mrc-3 - mrc-1* BC2 *aabb* | 6.379 | 2.695 | 53 | 2.367244 | 0.186556 | -1.588 | 14.346 |
| *mrc-1* BC2 *AABB - mrc-1* BC2 *aabb* | 9.787 | 2.695 | 53 | 3.632056 | 0.007919 | 1.820 | 17.754 |

D)

| Grain size (area mm²) | | | | | | | |
|---|---|---|---|---|---|---|---|
| contrast | difference | SE | df | t.ratio | p.value | - 95% CI | + 95 CI |
| WT - *mrc-1* | -0.684 | 0.668 | 53 | -1.02467 | 0.907566 | -2.658 | 1.290 |
| WT - *mrc-2* | 0.279 | 0.668 | 53 | 0.418244 | 0.998272 | -1.695 | 2.253 |
| WT - *mrc-3* | -1.951 | 0.668 | 53 | -2.92194 | 0.054466 | -3.925 | 0.0231 |
| WT - *mrc-1* BC2 *AABB* | -2.077 | 0.668 | 53 | -3.11028 | 0.033791 | -4.051 | -0.103 |
| WT - *mrc-1* BC2 *aabb* | -0.809 | 0.668 | 53 | -1.2119 | 0.82914 | -2.783 | 1.165 |
| *mrc-1 - mrc-2* | 0.964 | 0.650 | 53 | 1.482452 | 0.676692 | -0.958 | 2.885 |
| *mrc-1 - mrc-3* | -1.267 | 0.650 | 53 | -1.94926 | 0.384721 | -3.188 | 0.655 |
| *mrc-1 - mrc-1* BC2 *AABB* | -1.393 | 0.650 | 53 | -2.14276 | 0.281888 | -3.314 | 0.529 |
| *mrc-1 - mrc-1* BC2 *aabb* | -0.125 | 0.650 | 53 | -0.19236 | 0.999961 | -2.046 | 1.797 |
| *mrc-2 - mrc-3* | -2.230 | 0.650 | 53 | -3.43171 | 0.01411 | -4.152 | -0.309 |
| *mrc-2 - mrc-1* BC2 *AABB* | -2.356 | 0.650 | 53 | -3.62521 | 0.00808 | -4.278 | -0.435 |
| *mrc-2 - mrc-1* BC2 *aabb* | -1.089 | 0.650 | 53 | -1.67481 | 0.55407 | -3.010 | 0.833 |
| *mrc-3 - mrc-1* BC2 *AABB* | -0.126 | 0.650 | 53 | -0.1935 | 0.99996 | -2.047 | 1.796 |
| *mrc-3 - mrc-1* BC2 *aabb* | 1.142 | 0.650 | 53 | 1.756897 | 0.501633 | -0.780 | 3.063 |
| *mrc-1* BC2 *AABB - mrc-1* BC2 *aabb* | 1.268 | 0.650 | 53 | 1.950401 | 0.384065 | -0.654 | 3.189 |

**Supplemental Table 3. KASP markers for genotyping the wheat mutants.** All primer sequences are given 5' to 3'. The wild-type allele primers had the VIC/HEX tail (GAAGGTCGGAGTCAACGGATT) on the 5' ends, while the mutant allele primers had the FAM tail (GAAGGTGACCAAGTTCATGCT) on the 5' ends. The nucleotide(s) that discriminate the wild-type from the mutated base (for wild-type and mutant primers) or the homeologous SNPs (for common primers) are indicated in capital letters.

| Gene | Line and mutation | Wild-type allele | Mutant allele | Common |
|---|---|---|---|---|
| *Tt*MRC-A1 | Kronos3272 (Q258*) | agcaacagttagggagctgC | agcaacagttagggagctgT | cctcgattcatttgatctggcg |
| | Kronos598 (L289F) | ccatcctaaactctgcttctcaaG | ccatcctaaactctgcttctcaaA | gaagcaacagttaggGAGcT |
| | Kronos4681 (Q550*) | catacggctcagatgctcgC | catacggctcagatgctcgT | gcaagatcgccagtgagC |
| 6B pseudogene | Kronos4305 | ttgagaagcagagtttaggatgG | ttgagaagcagagtttaggatgA | acgtttgaagtcagtgataataccA |
| | Kronos3078 | aggattcagagctttctgatacaC | aggattcagagctttctgatacaT | tgaagctcaGcaatttcactgc |

**Supplemental Table 4. Summary of leaf starch quantification experiments.** ED = End of Day, EN = End of night. Non-valid sample = sample lost during experiment or omitted because technical replicates were too divergent. All experiments were done in the same CER, on a 16-h day, 8-h night cycle.

| Experiment | Date | Genotypes | Time | Number of biological reps per genotype planned | Total non-valid samples deleted | Total outliers deleted from final dataset | Final n |
|---|---|---|---|---|---|---|---|
| **1** | Oct-19 | WT, *mrc-1, mrc-2, mrc-3* | ED+EN | 6 | 3 | 0 | ED: 6,6,5,6; EN:6,6,4,6 |
| **2** | Dec-19 | WT, *mrc-1, mrc-2, mrc-3* | ED+EN | 6 | 5 | 0 | ED: 6,6,6,5; EN: 4,6,5,5 |
| **3** | Jun-20 | WT, mrc-1, mrc-2, mrc-3 | ED | 20 | 8 | 0 | 18,16,19,19 |
| **4** | Jan-21 | WT, *mrc-1, mrc-1* BC2 *AABB*, *mrc-1* BC2 *aabb* | ED | 15 | 10 | 1 | 15,12,10,12 |
| **5** | Feb-21 | WT, *mrc-1, mrc-1* BC2 *AABB*, *mrc-1* BC2 *aabb* | ED | 12 | 3 | 0 | 10,12,12,11 |
