## Supplemental File 1 for "The plastidial protein MRC promotes starch granule initiation in wheat leaves but delays B-type granule initiation in the endosperm"

**Supplemental File 1. *Aegilops speltoides* *MRC* gene model**. The genome of *Aegilops speltoides* isolate TS01 was searched using BLASTn, using the *TaMRC-A1* coding sequence as the query. A match was identified on Chromosome 6 in the region from 180007045 to 180009267, with 97.5% sequence identity. Sequences highlighted in yellow represent the two putative exons of MRC, red text represents the start and stop codons, blue text represents splice junctions.

>CM038146.1:180003045-180012267 Aegilops speltoides isolate TS01 chromosome 6, whole genome shotgun sequence

TAATGGCTCTCTCCACGAATGAAGAGGGGGTGGAGGGGTATAAATAGCCTCCACACAAAATCCAACCGTTACACACATTTGAGCAATCTCGGTGGGACCAAAGTGGACAACACGGTGAGACCGATAGTTCAAATAATGTGAACGTTGGGATTTTTGGTGGGACCGAAAATACAATCTCGGTTAGTCCGATTCGTGCAATGACAAGAGCATGCCAATTTGGTGGGACCGACTTCATCATCTCGGTGACACCGATTTGATGATGTAGGTTCTGAAGCTTGACAACTCATGTTTGGTGAGACCGAGTTCGTAATCGAAAATCCGAGTTTTCTAGGGTTTGGCTTATGGTGAGGTGGTTCATCTCGGTGTGATTGAGTTTGGAATATCGGTGGGACCGAGACTACTTATGAGTGTATATGGCGGTGAAAGTTCTTGATAGTTTAGGAGATGTATCACTAAGCACTTGAGCAACAGATCATCATCATAACCTCATCCCTTTTAATAGTATTGGCTTTTCCTATGGACTCAATGTGATCTTGGATCACTAAACCAAAAATGGAGTCTTGAGCTTTTGCCAATGCTTGCCTTAGCATTTTGGGGGTCCACTTCTTGATCCTTGCCATACCCATATCAATGAACTCTCTGAAATGATTCTCAAGTAACTATTAGTTCAATGATATATATGTTGTTATTAATTACCAAAACCACTGGGATTAGTTGCACTTTCGGTTTGTAAGTTGGTGCGTTCTAAACAAAGCGTCGTGGCGGGATCTACATGTGAAGCGAGTACATAGCTGCTTCGAAGCAGCAAATGAAGGAGTGCGGATGAAGGAGTTCATTTCGATCTAGGTGTCATACCTAGTGCATCGGGACCAATGAAGATCTTCGTGACAATCTTTGGTGCAATTGCCTTGGCAAAAGAATCCAGATTTTACAAGAGGACCAAGCACATCAAGAGACGCTTCAATTCCATTCGGGACCAAGTCCTAGTGGGAGACATAGAGATTTGCAAGATACATACGGATCTGAATGTTGCAGACCCGTTGACTAAGCCTCTCTCACGAGCAAAACATGATTAGCACCAAGACTCCATGGGTGTTAGAATCATTACTATGTAATCTAGATTATTGACTCTAGTGCAAGTGGGAGACTGAAGGAAATATGCCCTAGAGGCAATAATAAAGTTATTATTTATTTCCTCATATCATGATAAATGTTTATTATTCATGCTAGAATTGTATTAACCGAAAACATGATAGATGTGTGAATACGTAGACAAACATATAGTCACTAGTATGCCTCTACTTGACTAGCTCATTAATCAAAGATGGTTATGTTTCCTAACCATAGACATGTGTTGTCATTTGATTAATGGGATCACATCATTATGAGAATGATGTGATTGACATGACCCATTCCGTTAGCCTAGCACTTGATCGTTTAGTATATTGCTATTGCTTTCTTCATGACTTATACAAAGTTCCGCAACTATGAGATTGTGCAACTCCGTTTACGGAAGAACACTTTGTGTGCTACCGGACTCACAACGAATCGGGTGATTATAAAGGTGCTCTCTGGTGTTTAGAAGGTACATGTTGGGTTGGCATAATTCGAGATTAGGTTTTGTCACTCCGATTATCGGAGAGGTATCTCTGGGCCCTCTTGGTAATACTCATCACCTAAGCCTTGCAAGCATTGTAACTAATGAGTTAGTTATAAGATGATGTATTACGGAACGAGTAAAGAGACTTTCCGGGAACGAGATTGAACTAGGTATTGGATACCGACGATCGAATCTCGGGCAAGTAACATACCGATGACAAAGGGAACAACGTATGTTGTTATGCGGTTTGACCGATAAAGATCTTCGTAGAATATGTACGAACCAATATGGGCATCCAGGTCCCGCTATTGGTTATTGACCGAGAATGGTTCTAGGTCATGTCTACATAGTTCTCGAACCCGTAGGGTCCGCACACTTAACGTTACGATGACAGTTTTATTATGAGTTTACAAGTTTTGATGTACCTAAGTTTGTTCGGAGTCCCAGATGTGATCACGGACATGACGAGGAGTCTCGAAATGGTCGAGACATAAAGATTGATATATTGGACGACTATATTCGGACACCTAAGTGTTAGGGTGATTTTGAGAAAATCGGAGTTGGGAGGGTTACCGAACCCCCGGAGAGTATTGGGCCTTATGGGCCTTAGGGAAAGGAGAGAGGGCGGCCAAGATGGGCCGCGCGCCTCCCCTCCGGGTCCGAATTGACTAGAGGGGGTGGCGCTTCCTTTCCTTCCCCCTTCCCTCCTAGTAGGAGTAGGAAAGGAGGAGTCCTACTCCTACTAGGAGGAGGATTCTCCTCCTTGGCACGCCTAAGGGCCGGTGGCCTCCCCTTTGCTCCTTTATATCTCGTGGGCGGGGGGCATAGACACACAAGTTGATCTACGGATCGTTCCTTAGCCGTGTCGGTGCCCCTCCACCATATTCCACCTCGGTCATATTGTCGCGGAGTTTAGGCGAAGCCTGCGTCGGTAGAACATCATCATCGTCACCACGCCGTCGTCTTGTGGAACTCATCTCCGGAGCTTTCTTTGGATCGGAGGCCGAGATCGCCATCGAAATAACGTGTCTTTGAACTCGGAGGCCCGACGTTCGGTGCTTGGATCGGTCGGATCGTGAGGACGACGACTACATCAACCGCGTTGTGTTAACGCTTCGCTTACGGTTCACGAGTGCGTGGACGAACACTCTCCCCTCTCGTTGCTATGCCATCACCATGATCTTGCGTGTGCGTAGGAAATTTTTTGAAATTACTACGTTCCCCAACACCGACATAGAGGAGTCGGCCACCTCCATCCCAAGGATGCAATTTTTTTTTTTGCGGGTGAAGGATGCAGTTCTTCGATAAGAAGTGTTTTCCAATAGCATATGGATGTTGACCTTTTATCCTAGTATTCCATTTCGAGTCAAAATAGTCCCTAACATTATAACAATTTTCATGTGTTACAAACTCCTAACAATTTTCATGTGTTATATAAACTCATTTCTTTTAAGGCAAACAAAGTGTACCTTAAGTTTCATGTGTTATACTCATAAATTTGGAACAGAAAGAAAAAAGGTAGTAAGACGAGTGAACGGGGAAGAAAAGCAGTAGAAGGCAAACGACGCAGCTCTCTCTCACGCTTCTCCCGTGGTCGACGTTGCAGTCCACACGCGGGCACGCGGCTGGGCGCGCCGGTTCCACCACCTCATCTCCCGCACTCCCTCTGCCTCGTATCTCGTCGCCTTCCTCCACACCCCGCAGGAGCATTGCCAGCCGTCCGATCGCGCCCGGGCGGCGGTGGTTCCCTCTCCCCATGTTCCGCGGCC**ATG**CGCCTCTCCATAGGCTCCCCATCCCCGTCGCCGCCGGCGGCGGTGGCCGCCGCTCTCCGCAGCACATCCCCGTCGTGCCGTACCGCCAGTCAT**GT**GAGCGCCCGCTGATCTTTTCTTCCTTTTCTCATATCGCTGTTTCGTGGTACCACGCTGCTCACTGTTACATGGACTGCTCGCGTTCGTGTTTTCCCGATTCCGTGCCCGTCCACACGTGTTTGAAGTAGAAGGATACTAGATTTGGTGTCCTAATTCATGTTCTGCTAGTACTAGTACTTTTTTTAAAAAAACTTTTCTGGAATTGGTTCGATTGTGATAAATTCAGTAAACTGCACCTGGCTGAACAAATCTTGATTGGAGAACGGCCTATGAACTCAAAAAAATTATACTGAACAGATGAAATGTTTATGCAGAGGTATGCTTGAGATCAAATTTCATCGGTTATGATACTTCACCTTATATGACAGTGAATTTCTGAAGTTCAGTGTACTGTCTTTCAGTTCGTCGATTTACAACAATTTTTTACGTGCTTAGTTTGAGGAAAGGATATTCCTCAGATTGCTTCACTAGGTTGTGACCATTTCCTTATCCTAATATCCTACTTATGCATTGTTTCCTGCAACTCTCTC**AG**GTTATGTTCAGGCAGAAGCTGAGTTTTATGGTGGCATTTCAGACTCAGCATCTGAAATATGCTCCTCGCTTGATCAAATCAGCCGTAAAAGGTATTAGATCAAATACAACTGATGGTGATAATGGAACGACTGAGCCAGCTAGAGAGTTGCTGGAGCGGCTATTTGCGAAGACACAAAGTTTAGACACTGGTGCTTCTCATGATAGTGAACTGAGCGTGAGCATTGAGGTCCTGAAGTCTGAATTCGAGGGTGCCTTGTCTATCCTCAGAAACAAAGAGAGGGATCTTCGCAGCGCAGAGAAGAGGGTTTCCGATGATCGGATAAGATTGAGCAAGACGAAGCAGGATCTTGATCAGAGAGAGGAAGCGATCCGCAAAGCTTATGTAAGGCAACAAGGAATAGAGAAAGCACTGAAAAAGGCAAGTAGAGATCTGGCGTTGCGAGTGAAGCAGATCAGTAATCTGAAGCTTTTGGTTGAGGGGCAAGACAGGACTATTGCCAGTTCACAAGCTTTGCTTTCTCAGAAGGTAACTGAAGTGGAAAATCTCAAACGAGATATGTTCAAGAAGAACGAGGAAGCAGACCTGATGCGTTCAGAGATCAGGTCCAAAGAACAGTTGGTTCTTACAGCTAATCAAGCTATTGCGCAGCAAGAAGCAACAGTTAGGGAGCTGCAAAGTGAAATTAAAAGAAAGACGATCGATATCGCCAGATCAAATGAATCGAGGAAAACTAATGAAGAGAAACTGAAAGTTGCTGAACAGGAACTTGAGAAGCAGAGTTTAGGATGGTTAGCAGCACAACAAGAGTTAAAGGAACTTGCACAACTGGCATTCAAAGATACAGATGATATCAATGGTATTATCACTGACTTCAAACGTGTGAGGTCTCTGCTAGATGCTGTACGCTCTGAATTAATCTCTTCAAAAGATGCTTTCGCTTCCTCTCGCAGACAAATAGAAGATCAAGCGGTTCAGTTGCAGGAACAAGTACAGGAACTCGAGGACCAAAGGGTATTACTGATGTCTTACACCCATGATTTGGAGGCTGCTAAACTGGAGATTCAAGGGAAGACACAGGAGCTCAGTTACGCACAGTCTCGTTGTCATGAACTTGAATCACAGTTACTTCAGGAAAGGGAGAAGGTCGAGTCTCTAGAAGCCGAATTAGCCAAAGAAAAACAGAGCTTAGAACATAGAACTGAAGAAGTAGGCTTTCTTCAGAAGGAGCTTGTTCAGAAAGAAAATGAGTGCACCAAATCACAAGAACTTGTTAAAGTAAAAGAGTTTGAGCTGTTAGAAGCCAGACAGGAAGTCCAAGATATGAAGTTAAAGGTAGAGTCTATTCAATTGGCTGTTCAAGAAAAGGATTCAGAGCTTTCTGATACACAGAGCAGACTAACTGAAGTCAGCAGTGAAATTGCTGAGCTTCAGCAGTTGCTAAATAGCAAGAAGGATCAACTGCTTCAGGCTAGAACTGAATTAGATGATAAAGAGCAACATATAGAAACACTGGAGAGTGAGTTGGATAGCATACGGCTCAGATGCTCGCAAGCTGAATCCATGGTTCAAAGGATGGCTGATCTCACTGGCGATCTTGCTAGTTCCGTAAAAGCCGGAGAAATGGACATCTATACATTACTGGATGATGAAATTTCAAGCACTGGTACAGCCCTCGAGTCCAATTTGCATAAGCATAATCAACTGGAGGCTGACATAGAGATGTTAAGAGAATGCTTGCGGCATAAGGACATGGAGTTGAGAGCTGCTCATGAAGCACTTGATGCCAAAGATCACGAGCTGAAGGCAGTACTTAGAAAGTGGGATGTGAAGGAGCGGGAAGTACGTGAGTTAGAAGAGTTACCGGATCCCAGTGCCACAAATGAACTTGCTGGTTTTTCCAGTGAGACAACAGAGGACGGCATTGTAGGAGAGATGGAGCTCCCAGAGCTTCAAATTGAAGCTGTGGAGGTCGAAGCACTTGCTGCTACGACTGCATTGAGGAAGCTAGCGGATATGACTAAGGATTTCTTCAAACACGGCAAAGCTGATTCTGGTATTGACTTGGTTGCATCAGAGAGTCAGAAAATCAGTAAATGTGATCCTAAAATGGAAGTACACAAGAAGACGGATGTGATTCTTGAAGCTGAAAAAGAAATAGTTAGGCTCTTCTCATTGACAAAACAGATTGTCACTGATGACATAATAAACGATATTGAGGAA**TGA**TAGCTTCAAACTGAAGCATGTAGTCTTCCAATTCTATCAAGATAGCTTCAGAGTAGAGATATACCAGATTGATCTTTCGAACATTTATGGACAGTGATGTCGCCCAGAAGGATGAGATCTTCTCTGGTTGATATCACAACTGCCATTTTGAAAAAGGGTAACATGTTGAGCAGAAGCTGGTCATCTGATCCTTTGTTCTCCTTTTGTAATGTACCTCAAACTATTCCTCAGATCTTTGTTCAATGTGTTCCTCCTAAATATACATGGGAAGTTATTGATATGGCAATTGGTGCTGGTTTTCGTCATGCCAAAACTACTACTGAGATTGTGGTGAATGTTTCCATGTGCAGCAAAGTTGGTGCAGCGATGTTGCAGCGTGTTTGTTATTTTACTGTTTCATTGCAACTTCAAATAGTTTCAAATCCCAAAGCTACTAAAGTAGATGGCTATCAAGTACTCTCTCTTTTTTCTTCAAAAAAGGTTAAATACTCTTGTCTTTCCACTTGCTTTCGAAAGCTCCGTAGTTCAAATTTACACAGGATGCTTGAGCAATTTACACAGGATGCTTGAGCATCAAACTGTTAGAAACAATGGGCCTAGTCCATGTACAATTTCTGAAATCTCAAATAGAGACCCATAATTAAAGGGATAATTAGATTTATGCCCCTAGTTGTGTCTCACTCAGCTGTTTTACCCCTAATTTCCAAGAGCCACCGGCTCTCTCCAAGTCACTTCGCTCCTCTTATACTTTTGCCATTTGACCGTTTGATCTTCAGTTTGAAAACCTCATAACTAATTCATACTAAATCAAAAAAATGCAAATAAGATTTCAAAATGTTCGGAAAAACATCACCTATGTGTCAGTGTCATTTGCATTCATGAAAAAAGTGTTGGAAAGTGCACATCTGAGTTTTAGCTCTTATGATACCACCATGAATAGTAAAATGTAGAAAAAAAAAATCAAAAAATTCAAAAACAAATTTGGTGGCAAAGAATGACAAATGTTGTAAGTGCTTGCCAAGTTTTATCAGGGAATGGCATCCGTGGATGTCGTCGCAAAAAAAAATCAGCACTCCAAAATAGATTTTTTTTTGCCACGACTTTCACGAATGTCGTTCCCTCATGAAACTTGGCAAGCACTTAAAACATTTGTCATTCTTTACCACCAATTTTTTTGAATTTTTTTATTTTTTTTTAAATTATACTATTCATGGTGGTATCATAAGAGCTAAAACTCGGATGAGCACTTTCCAACACTTTTTCATGAGTGCAAATGACAATGGCATATAGGTGATGTTTTTCCGAACATTTTGTTATCTTATTTGCATTTTTTTATTTAGTATGAATTAGTTATGAATTTTTCAAACTGATGGTCAAAGGTCATAGGGCAAAAGCATAAGAGGAGCAAAGTGACTTGGATAGAACCGGTGACTTTTGGAAATTATGGGTAAACAACCGAGTGGGTCACAACCAGGGGCATAAATCTAATTATCCCATAATTAAATGGCAAGTGGTGGTGCTAAAGTTTAGAGGAAGTGTTTCACCGCTTTGTATAGTGGGTTCTCTTTTTTTTCGAGAAACTTCCAATCTATTCATTTTCAATCATGGCAGTACAACGAACACCAGAAATAAAAATAATTACATCCAGATTCGTAGACCACCTAATGACTACTACAAGCACTGAAGCGAGCCGAAGGCGCGCCACCATCATCGCCCCTCCCTTGTCGGAGCCGGACACAACTTGTTGTAGTAGACAGTCGGGAAGTCGTCGTGATAAGATCGAAAGGATCAACACACCAGAACAGCAACCGCCGCCGATGAAGACAAACTTAGATCGAAAGGATCCAACCTCCACACTAAGTGATGTGTTGAGAAGAGAAATAGAAGACCACACATGCGCTTGCCTTGCCTCGCCTGGCCGTGCACAACTTCCTTTTGTAGTTTAATTTTTTGATGTCTTGGCAGACAAGTTAATTATTTCTTGTGTGGTAAGTATATGACTTAGAAACCAAGTTGGTTTGAGATCGTGATCACGACATGATACCGCTTCTGGTCCGTTACCAGCCGCGGCAAAAGACACAACGAAAACATCTAGGGTTTTGTCACATCTTGCAACTTACACTGCCACAGTAGTCTATTCCATCCCGAACGCCAACGTGCATCGGCGTGCGCGAGAGGAGGTCTCTGGAAGCGTTCGTCCTTGCGATTTTGCACCGGGAGAGGACGAATTAGGTTTTTGGGAAGTGCTCTGCGTGACTGTCCACGTTCTTCATCACGAGTTGTCTTCTGTCCAAGTCGGGCAGCACTACTCATTGTCATCTACAACAACGTTAGCAAAAGATTGTCGTCAACATCATCATCAACAACATCGCTCCTGCAGTAGCTAACGAACAGTACATCCATTGTAATCTGTTCATGTCTCTATTTGTAGTTATTGTTACATATGTGATGATGCCGTGCATGTTATCTGGTTTGTCTAGTATGCTAGATTATTGCATGCTATTTTTTGTAGTACCATTTATGAATTATTTACTGGAATTAATATTGGATTTGCCTAATATTCCAACACAAACGTGTACAAGCTAAGAAACTTGAGGCTAGAGTTTAACTTTGAATCATCGAACCAACTATTTAAATCCATGGCAGACTAAGGCATAGCCTAGTGGTGGGAAGGGGCTGATGCCTTCCCCCCACCCAGATTCAAGGCATGGTACTTGCAAATTGGGTTTGTTGCACCAATTATACTGTAGGGGGTTCTCTTACAGTCTTTCAGTCAAAAAAAAAATATATTTAAATCCATGCCGAAGAAACTCATGGTCTTGATCATTCTAGCACGCATATAAGTGGTTAACAAAAAAGCACGTCGTCATCTCTTCGGAGACTATTGATATCGATAAATGTGACTCAAAAATGGAAGTACACAAGAACATGGTGATCAGACATTGTAGCCCCGTGTTGTCACCTCGACAAGGCGACAGTGACAACAGGGTTGATCAA
